# Evaluation of new adjuvants in enhancing the immune response to a candidate of Severe Fever with Thrombocytopenia Syndrome Virus vaccine

**DOI:** 10.64898/2026.09.11.750931

**Authors:** Jorge Levican, Natalia Ferressini Gerpe, Seok-Chan Park, Moataz Noureddine, Farah El-Ayache, Vivian Yan, Leonie Gruneberg, Mathew Prellberg, Lauren Chang, Gabriel Laghlali, Gagandeep Singh, Pamela T. Wong, Arturo Levican, Gustavo Palacíos, Adolfo García-Sastre, Michael Schotsaert

## Abstract

Severe fever with thrombocytopenia syndrome (SFTS) is an emerging zoonotic disease caused by the tick-borne bunyavirus SFTSV. SFTSV is primarily transmitted by *Haemaphysalis longicorn*is ticks and can cause a hemorrhagic fever-like illness characterized by high fever, thrombocytopenia, leukopenia, and multiorgan failure, with reported case-fatality rates ranging from 10% to over 30%. Despite its growing public health relevance, no licensed vaccines or specific antiviral therapies are currently available.

Here, we evaluated a beta-propiolactone-inactivated SFTSV vaccine candidate based on strain YL1, formulated with benchmark adjuvants (Addavax and Alhydrogel) or experimental innate immune agonists, including Sendai virus-derived defective interfering RNA (SDI), a RIG-I agonist, and imidazoquinoline-poly(ethylene glycol)-cholesterol conjugate (IMDQ-P-C), a TLR7/8 agonist, alone or in combination. Using an extended prime-boost regimen, we examined adjuvant-dependent modulation of humoral immunity, antigen-specific cellular recall, antibody cross-reactivity, and functional protection following passive serum transfer and live-virus challenge in IFNAR-/- mice.

All adjuvanted formulations induced detectable humoral responses, with Addavax and SDI eliciting the highest virus-specific binding and neutralizing antibody titers. Antigen-specific analyses identified the nucleoprotein (NP) as the dominant target of vaccine-induced immune memory. NP elicited robust antibody response and the broadest cytokine recall profile following *ex vivo* stimulation. This response was broader than that induced by the whole inactivated virus and was characterized by mixed Th1-associated, Th2-associated, and proinflammatory-associated features that varied across adjuvant formulations. In contrast, the head domain of the Gn glycoprotein elicited limited cellular recall responses, consistent with its relatively weak humoral immunogenicity after both priming and boosting. Nevertheless, sera from SDI and Addavax-adjuvanted groups displayed cross-reactivity with Gn from a heterologous SFTSV strain, suggesting broader antibody recognition across viral genotypes.

Functionally, sera from SDI and Addavax-immunized mice conferred partial protection following passive transfer into IFNAR-/- recipients, as reflected by reduced viral loads, delayed disease onset, and prolonged survival after lethal SFTSV challenge. Together, these findings support the immunogenicity and protective potential of a BPL-inactivated SFTSV vaccine platform and identify NP as a major target of vaccine-induced immune memory. They further highlight the importance of adjuvant selection in shaping the magnitude, antigenic focus, and functional quality of vaccine-induced antiviral immunity.

**In brief:** Severe fever with thrombocytopenia syndrome virus (SFTSV) is an emerging tick-borne bunyavirus that causes a hemorrhagic fever-like illness with high fatality rates and lacks approved vaccines. Here, we developed and characterized a β-propiolactone-inactivated whole-virus SFTSV vaccine candidate formulated with benchmark and experimental adjuvants, including the RIG-I agonist SDI and the TLR7/8 agonist IMDQ-P-C. In BALB/c mice, Addavax and SDI elicited the strongest virus-specific binding and neutralizing antibody responses. Antigen-specific recall analyses identified nucleoprotein (NP) as the dominant target of vaccine-induced immune memory, eliciting robust antibody responses and the broadest cytokine recall profile after ex vivo stimulation. NP-induced recall showed mixed Th1-associated, Th2-associated, and proinflammatory-associated features that varied across vaccine formulations, whereas the head domain of the Gn glycoprotein elicited limited humoral and cellular responses. Sera from Addavax- and SDI-adjuvanted groups displayed cross-reactivity with the Gn glycoprotein of the heterologous HB29 strain and conferred partial protection following passive transfer into IFNAR-/- mice challenged with live SFTSV. These findings identify NP as a major immunological target of this inactivated vaccine platform and highlight how adjuvant selection shapes the magnitude, antigenic focus, and functional quality of antiviral immunity.

## Introduction

The ongoing transformation of the global climate is exerting profound effects on ecosystems and public health. Rising temperatures, altered precipitation and humidity patterns, and ecosystem disruptions have collectively expanded the geographical distribution and seasonal activity of key arthropod vectors, including ticks and mosquitoes (De Souza & Weaver, 2024). In parallel, increasing anthropogenic encroachment into previously undisturbed habitats has intensified the frequency and proximity of interactions between humans, domestic animals, and wildlife. As a result, regions once unaffected by specific pathogens are now experiencing the emergence and re-emergence of vector-borne infectious diseases, reshaping the global landscape of epidemiological risk (De Souza & Weaver, 2024) .

In this context, a new acute infectious illness, with hemorrhagic manifestations, was described in 2009 in rural settings in Hubei and Henan in central China, which was identified as the severe fever with thrombocytopenia syndrome (SFTS). Although the clinical presentation was initially thought to resemble human anaplasmosis, a new virus was identified by viral culture, electron microscopy, and nucleic acid sequencing (X.-J. Yu et al., 2011). The virus was provisionally named Severe Fever with Thrombocytopenia Syndrome Virus (SFTSV) and, according to the latest ICVT classification, it is currently named *Bandavirus dabieense,* belonging to the *<u>Bandavirus</u>* genus and *<u>Phenuiviridae</u>* family (*Taxon Details | ICTV*, n.d.; X.-J. Yu et al., 2011). Since its discovery, SFTSV has been detected in several East Asian countries, including South Korea, Japan, and Vietnam, and is now considered a major emerging life-threatening disease (Cui et al., 2024; He et al., 2020; Tran et al., 2019).

The illness is characterized by thrombocytopenia, leukocytopenia, gastrointestinal disturbances, and hemorrhagic manifestations. In severe cases, the disease can progress to disseminated intravascular coagulation, multi-organ failure, and death (Wang et al., 2021). Reported case-fatality rates range from 10% to over 30%, and no vaccine or specific antiviral treatment is available, underscoring the urgent need for effective preventive measures in endemic areas (Cui et al., 2024; He et al., 2020; Y. R. Kim et al., 2018; Miao et al., 2021; X. Wang et al., 2021).

The viral cycle includes wild and domestic animals as reservoirs and is transmitted to humans through the bite of its arthropod vector, the tick *Haemaphysalis longicornis*. Currently, growing evidence indicates that human-to-human transmission may also occur, particularly in healthcare settings (Hu et al., 2022; Y.-X. Wu et al., 2022, 2022; N. Zhang et al., 2024).

SFTSV is a single-stranded, negative-sense, tri-segmented RNA virus. Its genome includes the small (S) segment that encodes the nucleocapsid protein (NP) and a non-structural protein (NSs), the medium (M) segment encodes the envelope glycoproteins, glycoprotein N (Gn) and glycoprotein C (Gc), and the large (L) segment encodes the RNA-dependent RNA polymerase (RdRp).

The nucleocapsid protein NP is the more abundant structural component of the virion and is an important focus of the immune response (L. Yu et al., 2012). NP resides in the viral core, enwrapping each genomic viral RNA (vRNA) to form the ribonucleoprotein (RNP) complex, which is essential for RNA replication and transcription. The non-structural protein (NS) of SFTSV plays a significant role in innate immune evasion by antagonizing the host interferon response, which is crucial for the early control of viral infection (Khalil et al., 2021). The surface glycoproteins Gn and Gc assemble in heterodimers that organize in hexons and pentons to form the icosahedral viral shell of approximately 110 nm in diameter (Du et al., 2023; Z. Sun et al., 2023; P. Wang et al., 2020). These glycoproteins also play critical roles in viral entry. The Gn glycoprotein mediates attachment to the host cell via the C-C motif chemokine receptor 2 (CCR2), while the Gc glycoprotein facilitates the fusion of viral and endosomal membranes, enabling ribonucleoprotein complex release into the cytoplasm (Spiegel et al., 2016; Tani et al., 2016; Y. Wu et al., 2017; F. Yuan & Zheng, 2017; L. Zhang et al., 2023). As the most outwardly exposed elements, these two glycoproteins represent a significant target of the immune response and are the focus of neutralizing antibodies (Liu et al., 2025; Moming et al., 2021; Sun et al., 2023). In addition, both glycoproteins are key targets for the development of therapeutic drugs, antibodies, and the rational design of vaccines (Dong et al., 2019; D. Kim et al., 2024; Kwak et al., 2019; S. Yuan et al., 2019; W. Zhang et al., 2020a, 2020b; X. Zhang et al., 2025a). Finally, the RdRp is responsible for viral genome transcription. Using the host-capped mRNA as a primer through the cap-snatching mechanism, the RdRp synthesizes an early positive-sense complementary RNA (cRNA) that serves as a template to produce new genomes during the viral replication (Reguera et al., 2016).

The emergence of SFTS and SFTSV is not an isolated phenomenon; instead, it reflects a broader trend of emerging tick-borne viral diseases. A notable parallel is the Heartland virus, first identified in the United States in Missouri in 2009. Like SFTSV, it is transmitted by a tick, the lone star tick *Amblyomma americanum*, and causes a similar clinical syndrome characterized by fever, leukopenia, and thrombocytopenia (Esguerra, 2016). Given the increasing incidence of this group of diseases and the potential for outbreaks, ongoing research into the biology of these viruses and their pathogenesis is critical for public health preparedness.

Although several SFTSV vaccine platforms have been described, the magnitude, antigenic focus, durability, and functional balance of vaccine-induced immune responses remain only partially characterized. In addition, prolonged prime-boost intervals have not been systematically evaluated for SFTSV vaccine candidates. Current evidence from other vaccine platforms suggests that extending the interval between priming and boosting can favor immune maturation and enhance the quality, durability, and recall potential of vaccine-induced responses. For example, a 16-week interval between doses of SARS-CoV-2 mRNA vaccines enhanced the magnitude and maturation of the antigen-specific B cell compartment, although its effect on T cell immunity was more limited (Nicolas et al., 2023). Similarly, in an H56/alum mouse model, an 18-week prime-boost interval increased germinal-center B cells and antibody-secreting cells while modulating the functional profile of reactivated CD4 T cells, despite not significantly increasing serum IgG titers (Pettini et al., 2021). Delayed boosting has also been associated with increased magnitude and durability of antigen-specific B cell and IgG1 responses in the RH5.1/AS01B malaria vaccine platform (Nielsen et al., 2023). More recent studies further indicate that dose spacing can influence the functional quality of T cell responses and inflammatory or reactogenicity-associated signatures, although these effects appear to depend on the vaccine platform and regimen used, particularly in SARS-CoV-2 mRNA and vector-based vaccines (Ahmed et al., 2025; Amini et al., 2025; Murray et al., 2025). These findings support the rationale that extended prime-boost regimens may provide a favorable immunological window for the maturation, persistence, and functional diversification of vaccine-induced immunity.

To address this gap, we evaluated a beta-propiolactone-inactivated SFTSV vaccine candidate based on strain YL1, formulated with benchmark adjuvants (Addavax and Alhydrogel) or experimental innate immune agonists, including Sendai virus-derived defective interfering RNA (SDI), a potent RIG-I agonist, and imidazoquinoline-poly(ethylene glycol)-cholesterol conjugate (IMDQ-P-C), a TLR7/8 agonist, alone or in combination (Jangra, De Vrieze, et al., 2021; Jangra et al., 2022; Martínez-Gil et al., 2013). Using an extended prime-boost regimen, we determine how adjuvants influence the magnitude and quality of vaccine-induced immune responses. Mice received a booster immunization 21 weeks after priming, providing an exploratory framework to assess immune persistence, post-boost antigen-specific cellular recall, and the protective capacity of vaccine-induced antibodies following passive transfer and live-virus challenge. Through this approach, we aimed to explore their ability to modulate the immune response and enhance protective efficacy, contributing to the rational design of improved vaccine strategies against this emerging pathogen.

## Materials and Methods

### Virus strains

The SFTSV virus, strains YL1 and HB29, and the Heartland virus strain MO4 were obtained from the University of Texas Medical Branch Arbovirus Reference Collection. The strain YL1 was isolated in China in 2011 and had 3 Vero cell passages. The HB29 was isolated in China in 2012 and had a history of 10 passages in Vero cells, plus one passage in SM1, and an additional passage in Vero cells. The HRTV MO4 strain has one passage in DH82 cells, one in Vero cells, followed by one in SM1 cells. The viruses were propagated in VERO E6 cells, titrated, and the strain selected for the vaccine formulation, YL1, was fully characterized by complete genome sequencing.

### Cells

Vero-E6 cells were maintained in DMEM supplemented with 10% FBS (unless another percentage was stated). For virus production, serum-free medium was used to avoid contamination with proteins of animal origin. VERO E6 cells were seeded at high confluence (60-70%) in OptiMEM medium supplemented with GlutaMax I. HEK293 cells were maintained in DMEM supplemented with 10% FBS (unless another percentage was stated).

### Virus titration

TCID50 was conducted in Vero E6 cells. Cells were seeded in a 96-well plate at a density of 50,000 cells per well in 200 microliters of 2% FBS DMEM. Cells were incubated overnight at 37°C with 5% CO2. Infection was performed using 200 microliters of a 10-fold dilution of the virus, and the infection proceeded for 4-7 days, for visualization by indirect immunofluorescence for SFTSV YL1 and HB29 or until the cytopathic effect (CPE) was visible for HRTV. The TCID50 was calculated using the Reed-Muench method (Reed & Muench, 1938) **Recombinant proteins.** The recombinant nucleoprotein NP was produced using ChampionTM pET Directional TOPO® Expression Kit (ThermoFisher). The complete N ORF from the strain SFTSV YL1 or SFTSV HB29 (nt 43-780 segment S, reference sequence: HM745932) was cloned downstream of the 6xHis-Tag in the vector PET151/D-TOPO. The resulting constructs were transformed into the expression-competent host BL21 StarTM (DE3) One Shot Chemically Competent E. coli kit (Thermo Fisher), under the induction of 0.5 mM Isopropyl β-D-1-thiogalactopyranoside (IPTG) for 12 hours at 25 °C. The nucleoprotein was further purified using the gravity flow Immobilized Metal Chelate Affinity Chromatography (Qiagen).

The complete ectodomain or the head domain of the glycoprotein Gn and the complete ectodomain of the glycoprotein Gn were cloned into the pCAGGS expression plasmid and produced in the mammalian cell Expi293F according to the manufacturer’s instructions (Thermo Fisher) and purified using gravity flow Immobilized Metal Chelate Affinity Chromatography (Qiagen). All the recombinant proteins produced were confirmed by SDS-PAGE and western blot using anti-HisTag secondary antibody (Supplementary Figure 1C).

### RTqPCR

RNA extraction was conducted using Trizol (Thermo Fisher), following the manufacturer’s instructions. The quantification of SFTSV viral RNA was performed following the protocol previously described (Perez-Sautu et al., 2020; Y. Sun et al., 2012). The quantification of the genome copy was determined by interpolating the sample CT value in a standard curve derived from serial dilution of a synthetic RNA fragment coding a partial S gene segment, produced by T7-dependent *in vitro* transcription.

### Cell transfection and Gc glycoprotein expression

HEK293 cells were seeded in a 48-well plate pre-coated with Ply-L-Lysine and allowed to attach overnight. The next day, cells were transfected with expression plasmids containing the complete open reading frame of the glycoproteins Gc and Gn, and nucleoprotein N of SFTSV, as well as GFP or an empty expression plasmid, using Lipofectamine 3000. A total of 400ng of DNA was used per well. After 24 hours of incubation, the transfection was assessed by GFP expression, and cells were fixed using PFA 4% for 15 minutes, permeabilized with PBS-Triton 0.01% for 10 minutes, and blocked with PBS-BSA 2%. Mice serum from day 14 after immunization was used (dilution 1:10) for 2 hours to detect the SFTSV proteins. AlexaFluor 488 anti-mouse secondary antibody was used at a 1:500 dilution, and DAPI was used at a 1:2000 dilution for nuclear staining. As a positive control, a commercial rabbit polyclonal anti-Gc primary antibody (Abnova, PAB27173) was used at a dilution of 1:1000 and detected with anti-rabbit antibody AlexaFluor 488 conjugated.

### Virus inactivation and antigen preparation

The virus SFTSV strain Y1 was grown in serum-free medium. Five T175 flasks were seeded with VERO E6 cells to obtain 70-80% confluence on the day of infection. The infection proceeded at a multiplicity of infection (MOI) of 0.01, with 1-hour adsorption involving gentle agitation every 15 minutes. We adjusted the culture medium to 30 mL of OptiPRO SFM (Thermo Fisher), while maintaining the original inoculum volume. After 7 days of incubation, the supernatants were clarified by centrifugation for 10 minutes at 3000 × g and subjected to precipitation with polyethylene glycol (PEG; Abcam, Fisher Scientific) according to the manufacturer’s protocol. Four independent batches of virus production were produced. The virus inactivation was carried out using 0.05% beta-propiolactone (BPL) and incubation at 4 °C overnight with gentle mixing. The viability of the inactivated virus was tested using TCID50, and the product was fractionated and stored at -80°C. The vaccine was visualized in denaturing SDS-PAGE, and the total protein concentration was measured by Bradford assay.

### Adjuvant formulations and vaccination

The <u>S</u>endai virus <u>D</u>efective Interfering RNA (SDI) was composed of the Sendai virus defective interfering RNA segment was synthesized by *in vitro* transcription using HiScribe T7 High Yield RNA Synthesis Kit (New England Biolabs), followed by DNase I treatment using TURBO DNA-free kit (Thermo-Fisher) and purification with RNeasy purification kit (Qiagen) (Patel et al., 2013). The IMDQ-P-C was obtained from the conjugation of imidazoquinoline 1-(4- (aminomethyl)benzyl)-2-butyl-1H-imidazo[4,5-c]quinolin-4-amine (IMDQ)[25] with cholesteryl-poly (ethylene glycol) (PEG-Chol), yielding IMDQ-P-C (Jangra, De Vrieze, et al., 2021). The Addavax (similar to MF59) and Alhydrogel adjuvants were obtained from Invivogen. The vaccination was administered to 6- to 8-week-old female BALB/c mice (groups of 5) obtained from Jackson Laboratories, CT. Mice were housed with free access to food and water in a pathogen-free facility at the Icahn School of Medicine at Mount Sinai, New York. All mice received the vaccine via intramuscular injection of 50 μL into the hamstring muscles of the left hind leg.

The mice were immunized with 5 μg of beta-propiolactone-inactivated SFTSV as antigen for both prime and boost. The adjuvants were administered as follows: SDI (1 mg of SDI with 40 μg nanoemulsion, 1:40 ratio), IMDQ-P-C (100 μg), Addavax (50%), Alhydrogel (50%), or nonadjuvanted (PBS). All these formulations have been previously proven in our laboratory (Jangra, De Vrieze, et al., 2021; Jangra et al., 2022, 2024). Blood samples were obtained from all mice after the prime and after the boost. Seven weeks after the boost, mice were subjected to terminal intracardiac blood collection under deep anesthesia using isoflurane 5%. Spleen harvest was conducted after the mice died.

### Passive transfer

Male and female IFNAR-/- mice were obtained from Jackson Laboratories. The passive transfer was performed via intraperitoneal injection of 250 μL of pooled sera (6 mice per group, 3 male and 3 female). Each group received a sera pool from a unique vaccine formulation donor. A no-serum control was also included in this protocol. Mice were challenged with 5 TCID_50_ of the live SFTSV strain YL1 under animal BSL3 conditions. Blood samples were obtained 5 days post-challenge to check the transferred antibody levels and viral load. Body weight and general condition of mice were monitored up to 14 days after challenge. All animal protocols were conducted under the guidelines approved by the Icahn School of Medicine at Mount Sinai Institutional Animal Care and Use Committee (IPROTO202400000092).

### Enzyme-linked immunosorbent assay (ELISA)

Ninety-six well plates were coated with 50 μL of the diluted recombinant protein or whole virus at a concentration of 2 μg/mL and incubated overnight at 4 °C. The plates were blocked with PBS-T+3% (w/v) non-fat milk powder for 1 hour at room temperature. After removing the blocking solution, serial dilutions of the sera were made in blocking solution and incubated for 1 hour at room temperature. After three washes with PBS-T, the plates were incubated with 50 μL of secondary anti-mouse antibody conjugated with HRP for 1 hour at room temperature. After three washes, 100 µL of Sigma Fast OPD (Sigma) was added to each well and incubated for 10 minutes at room temperature. The reaction was stopped with 50 µL of 3 M HCl, and the absorbance at 490 nm was measured.

### Microneutralization assay

The neutralization activity of the sera from mice was tested post-prime and post-boost in all groups tested. Serial 2-fold dilutions of the sera were co-incubated with 100 TCID50s for 30 minutes. Then, the media of Vero E6 cells in 96-well plates was replaced by the virus-serum mixture, and the plates were incubated for 72 hours for SFTSV and 4 days for HRTLV. Virus alone, serum alone, and only media were used as positive and negative controls. Back titrations of the viral stock were performed simultaneously.

Indirect immunofluorescence was used as a readout of SFTSV infection. Immunostaining was performed using a rabbit polyclonal anti-SFTSV antibody (Abnova) and a goat anti-rabbit Alexa Fluor 488-conjugated secondary antibody (Thermo Fisher). To check for HRLV infection, plaque-forming assays were used as a readout, staining the cell culture wells with crystal violet after paraformaldehyde (7%) fixation.

### T cell antigen recall assay

The T cell antigen recall response was conducted using splenocytes isolated from immunized mice 7 weeks after the boost as described previously (Jangra, Landers, et al., 2021). Briefly, isolated cells were plated at a density of 8×10^5 cells/well and stimulated with 5 μg per well of whole inactivated virus (YL1 strain), the recombinant N nucleoprotein or the recombinant head domain of the glycoprotein Gn in T cell media (DMEM, 5% FBS, 2 mM L-glutamine, 1% NEAA, 1 mM sodium pyruvate, 10 mM MOPS, 50 μM 2-mercaptoethanol, 100 IU penicillin, and 100 μg/mL streptomycin), in a total volume of 200 μL per well. Cells were stimulated for 72 h at 37°C, and secreted cytokines (IFN-γ, IL-1β, IL-2, IL-4, IL-5, IL-6, IL-12p70, IL-13, TNF-α, and IL-18) were measured in supernatants using ProcartaPlex Multiplex Immunoassay (Invitrogen, ThermoFisher). Cytokine responses were expressed as the antigen-induced change in cytokine concentration relative to the corresponding unstimulated condition (Δ concentration). For each cytokine, the concentration measured in the unstimulated condition was subtracted from the concentration measured after antigen stimulation. Before this calculation, negative concentration values of small magnitude and values below the assay limit of detection (LOD) were replaced with the corresponding LOD as appropriate. Negative Δ values were retained and were interpreted as decreased cytokine expression following antigenic stimulation relative to the matched unstimulated condition.

### Statistical analyses and data visualization

To evaluate the normal distribution, the Shapiro–Wilk test was applied. Parametric (one-way ANOVA, unpaired t-test) or non-parametric tests (Kruskal–Wallis, Dunn’s multiple comparison) were applied as appropriate. A p-value < 0.05 was considered statistically significant. Arithmetic or geometric means were used for normally distributed variables, while the median and interquartile range (IQR) were used for non-parametric variables. Those analyses and the graphical representation were computed using GraphPad Prism 10 for macOS, Version 10.5.0 (673). The cytokine analyses concentrations were expressed as Δ relative to the unstimulated control group using the following formula: Δ concentration = [Adjuvanted vaccine Stimulated condition – [Adjuvanted vaccine Unstimulated condition]. For the global antigen-specific recall analysis, cytokine responses expressed as delta cytokine concentrations were tested against zero using two-sided one-sample Wilcoxon signed-rank tests. To account for multiple testing, p-values from the 30 antigen-cytokine comparisons were adjusted using the Benjamini–Hochberg procedure to control the false discovery rate. Benjamini–Hochberg-adjusted p-values < 0.05 were considered statistically significant.

## Results

To prepare the antigen for vaccine production and minimize the presence of host-derived proteins, VERO E6 cells were adapted to grow in serum-free medium. Under these conditions, the virus displayed robust replication kinetics as determined by genome copy equivalents per μL quantification (Supplementary Figure 1A) and reached high infectious titers, exceeding 10^7^ TCID /mL, which were suitable for downstream inactivation and formulation (data not shown).

Polyethylene glycol (PEG) has been a widely utilized reagent for concentrating diverse viruses from liquid samples due to its ability to induce precipitation through volume exclusion and macromolecular crowding (Deboosere et al., 2011; Flood et al., 2021; McLellan et al., 2021). Using this technique, we achieved a 295-fold concentration of our initial viral suspension, as determined by TCID_50_ (data not shown). This viral concentrate was inactivated using 0.05% beta propiolactone (BPL), and the viral viability test showed no remaining viability of the viral stock. Finally, the PEG-concentrated and BPL-inactivated virus was analyzed by denaturing SDS-PAGE (Supplementary Figure 1B), followed by protein quantification, aliquoting, and storage at -80°C.

### 1. Vaccine immunogenicity

To check the immunogenicity of the inactivated virus, we conducted a pilot experiment using intramuscular immunization with 10 μg of BPL inactivated virus adjuvanted with Addavax and a nonadjuvanted condition (PBS) in BALB/c mice. After 14 days, the mice were bled, and serum reactivity was tested in ELISA. The humoral response was assessed by ELISA using the whole virus and the recombinant nucleoprotein NP, the full ectodomain and the head domain of the Gn glycoprotein, and the full ectodomain of the Gc glycoprotein as coating antigens (Figure 1A).

**Figure 1.**
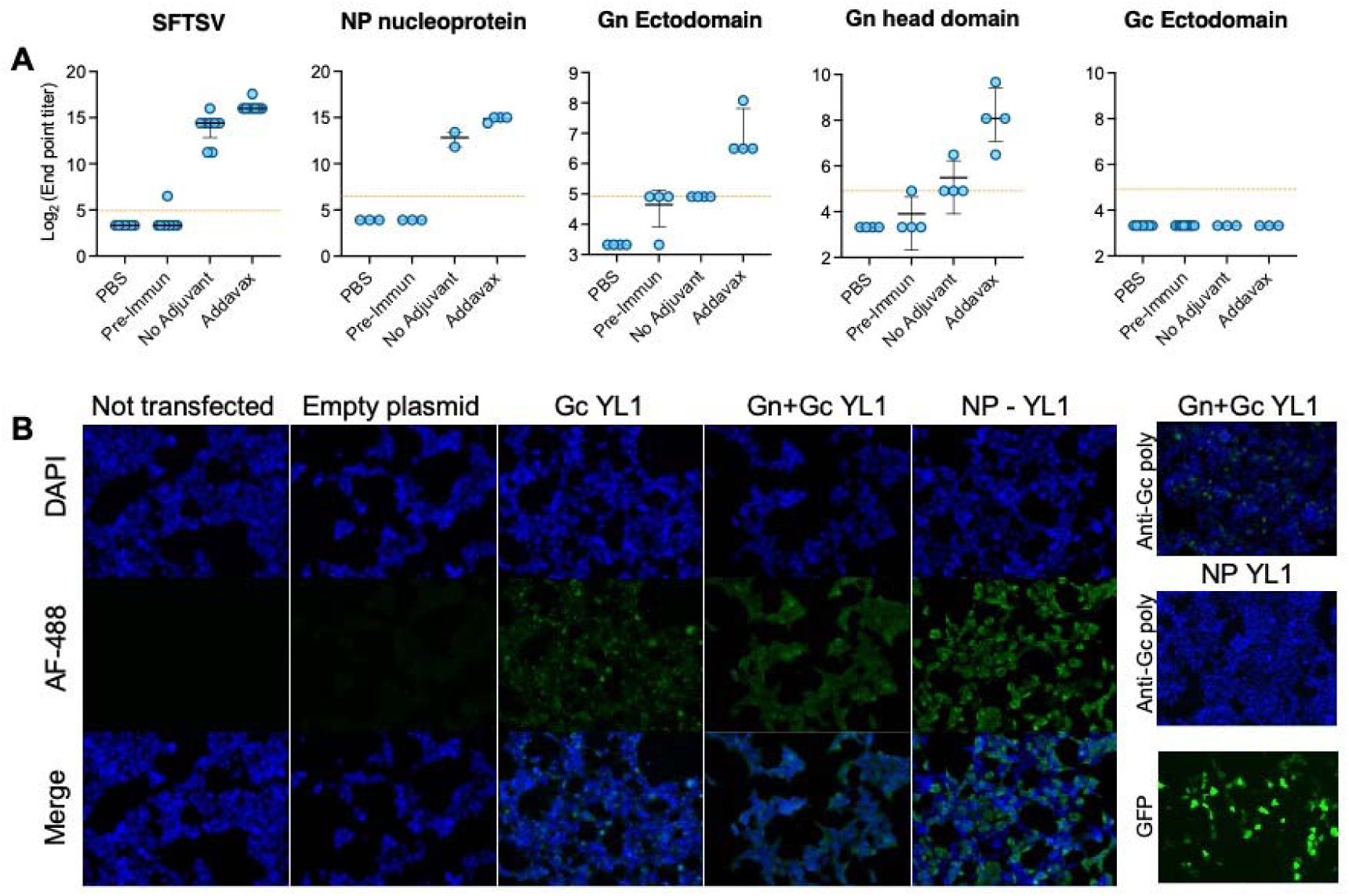
SFTSV-BPL inactivated Vaccine immunogenicity evaluation. **A.** Antibody response was evaluated 2 weeks after immunization with 10 μg of BPL-inactivated SFTSV vaccine, with and without adjuvant (Addavax). The humoral response directed to the whole virus (same as the vaccine) and its structural components: the nucleoprotein N, the glycoprotein Gn (both the complete ectodomain and the head domain), and the ectodomain of the glycoprotein Gc, was evaluated. The graphs display the median and interquartile range (IQR) of the individual titers, defined as the reciprocal of the last positive dilution (end point titer). The dotted line shows the limit of detection (32), and the dots representing non-reactive sera were plotted below this line for graphical representation only. **B.** The complete ORF of the glycoprotein Gc was expressed in HEK293 cells and tested with vaccinated mouse sera to check the presence of anti-Gc antibodies. Glycoprotein Gn and nucleoprotein N (both full ORF) were also expressed as controls. A commercial anti-Gc antibody was used as a positive control, and a GFP-coding expression plasmid was used to assess the transfection performance. Empty plasmid transfection and non-transfected controls are also shown. After 24 hours of transfection, the expression of GFP was evaluated, and the transfected cells were incubated with the respective sera or control. The reactivity was then revealed by indirect immunofluorescence using an Alexa Fluor 488 anti-mouse secondary antibody and DAPI nuclear staining.

This analysis shows high antibody titers against the full inactivated virus and the nucleoprotein NP. In addition, a modest reactivity was observed against the glycoprotein Gn, both the complete ectodomain and the head domain, whereas no response was observed for the glycoprotein Gc (Figure 1A). Although not statistically significant, a higher humoral response was observed in the adjuvanted formulation compared to the non-adjuvanted formulation.

To assess whether the lack of reactivity with Gc was due to poor immunogenicity or to the absence of native epitopes resulting from structural misfolding of our expressed glycoprotein, we expressed full-length Gc, with or without Gn, in HEK293 cells. Using a commercial polyclonal antibody against Gc, we confirmed its expression and reticular localization (round peri-nuclear pattern, upper right Gn+Gc YL1 in Figure 1B). Importantly, sera from mice immunized with the inactivated virus recognized the full-length Gc in transfected cells, but not the truncated ectodomain used in ELISA (panel Gc YL1 in Figure 1B, negative control panel in Supplementary Figure 1D). This observation suggests that the recombinant Gc may lack conformational fidelity. Consequently, Gc was excluded from subsequent humoral analyses, which focused on the whole virus, NP, and the Gn head domain.

### 2. Vaccine formulations and immunization schedule

To evaluate the immunogenicity of the inactivated SFTSV vaccine platform and the contribution of distinct adjuvants, 8-week-old female BALB/c mice (n=5 per group) were immunized intramuscularly using a homologous prime-boost regimen. Mice received 5 μg of β-propiolactone-inactivated SFTSV formulated with Alhydrogel, Addavax, SDI, IMDQ-P-C, IMDQ-P-C plus SDI, or without adjuvant.

We used and extended a prime-boost interval schedule to assess the persistence of the primary vaccine-induced response and its capacity for recall after a prolonged resting period. At 18 weeks after priming, serum samples were collected to evaluate antigen-specific antibody responses against whole SFTSV, nucleoprotein NP, and the Gn head domain by ELISA. Serum neutralizing activity was also assessed at this time point.

At week 21 post-prime, mice received a homologous booster dose containing 5 μg of β-propiolactone-inactivated SFTSV formulated with the corresponding adjuvant or without adjuvant. Three weeks later, serum samples were collected to reassess antigen-specific binding and neutralizing antibody responses. At seven weeks post-boost, mice were euthanized, and terminal blood samples were collected by intracardiac puncture for subsequent passive serum-transfer experiments. Spleens were harvested simultaneously for *ex vivo* T cell-antigen recall assays (Figure 2A). This schedule enabled the evaluation of antigen-specific antibody persistence after priming and the assessment of humoral and cellular recall responses following a prolonged interval between antigen exposures (Figure 2A).

**Figure 2.**
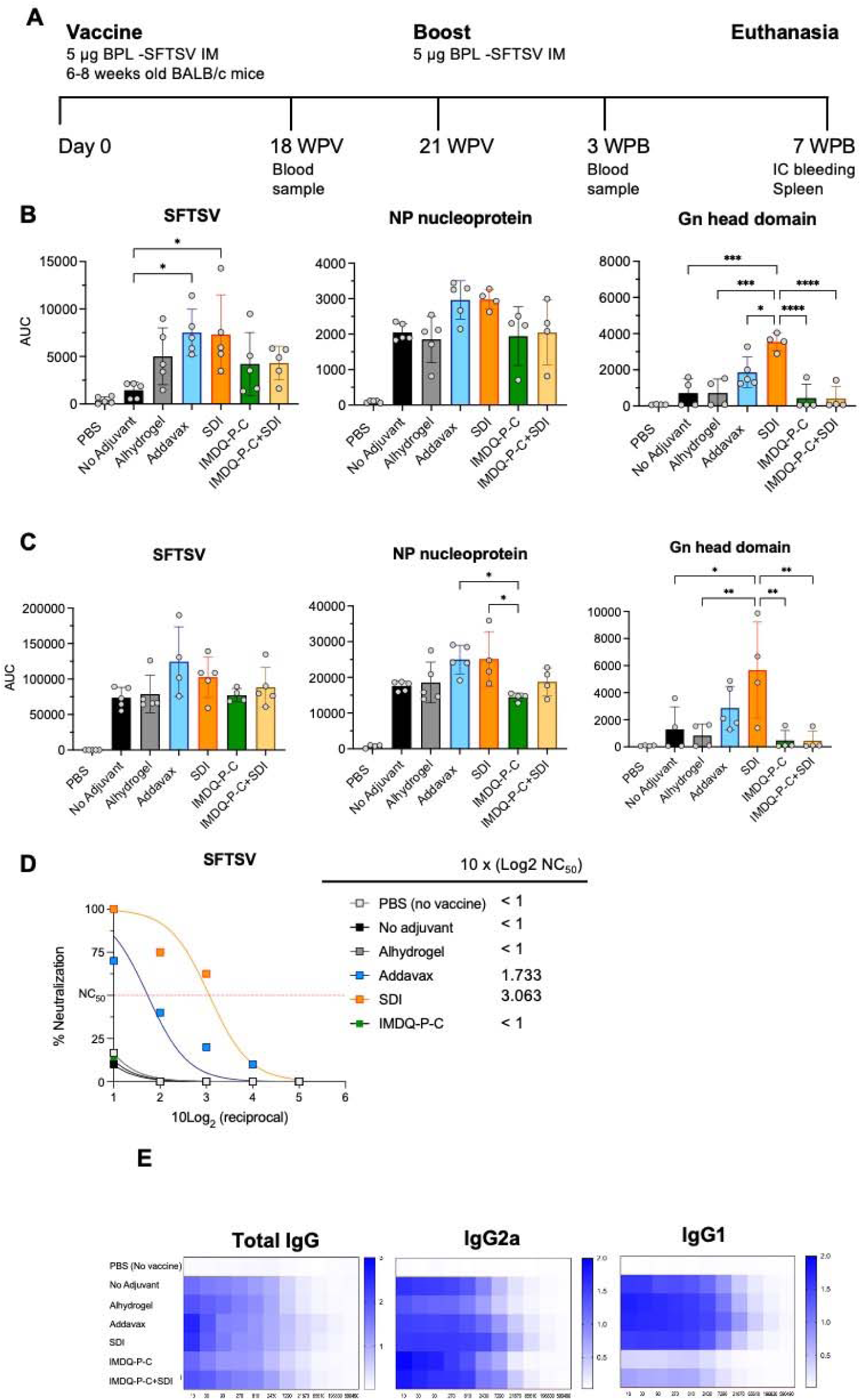
Vaccination scheme and humoral response after priming and boost. **A.** The priming and boost scheme is illustrated, including the sampling points for blood testing to assess humoral response, the terminal intracardiac blood collection, and spleen harvest. **B.** Humoral response after priming. The Mice were immunized with the BPL-SFTSV vaccine using different adjuvanted or nonadjuvanted formulations, as well as only PBS (No vaccine), and the humoral response was evaluated by ELISA using the whole virus (SFTSV), the nucleoprotein N, and the head domain of the glycoprotein Gn. The area under the curve (AUC) of the reactivity of each serum was determined, and the mean and standard deviation are shown. The statistical difference within groups was determined by one-way ANOVA, and the paired differences were computed using Tukey’s correction for multiple comparisons. P<0.05 was considered statistically significant. Paired comparisons between all adjuvants and PBS (no vaccine) were statistically significant, although the brackets and asterisks are not shown for image clarity. **C.** Humoral response after boost. Three weeks post-boost, the humoral responses were quantified and analyzed as in B. The mean and standard deviation of the antibody responses are shown for each condition (p < 0.05). **D.** Neutralization assay was performed using serial twofold dilutions of sera from 3 to 5 mice per group. The tests were conducted in duplicate. Neutralization efficacy was expressed as the percentage of wells neutralized per group, and non-linear regression analysis was performed using the least squares regression fitting method. The neutralization concentration of 50 was calculated, and the values are displayed as 10Log2 of the reciprocal dilution. **E.** Antibody reactivity was assessed by ELISA using whole inactivated SFTSV as the antigen, with subtype differentiation performed using anti-IgG1 and anti-IgG2a secondary antibodies. Three heatmaps are shown: total IgG, IgG1, and IgG2a responses. Reactivity is depicted as optical density (OD) values at 492 nm across serial reciprocal dilutions of mouse sera. The corresponding color bar is shown alongside each heatmap.

### 3. Persistence of humoral response after priming

In line with the response at 14 days post-priming observed in the pilot experiment, at 18 weeks post-priming, antigen-specific binding antibody responses against whole SFTSV and nucleoprotein NP remained detectable across vaccinated groups (Figure 2B). Addavax and SDI-adjuvanted formulations induced the highest reactivity against whole SFTSV, with both groups showing significantly higher responses than the non-adjuvanted formulation (p < 0.05). These formulations also showed higher reactivity against nucleoprotein NP than the other groups, although these differences did not reach statistical significance.

Differences between adjuvant formulations were more apparent when reactivity against the Gn head domain was assessed. Addavax and SDI were the only formulations that induced consistent Gn-specific antibody responses in all mice within their respective groups. In particular, SDI induced significantly higher Gn-specific reactivity than all other formulations (p < 0.05; Figure 2B).

The viral glycoprotein Gn is recognized as a major target for neutralizing antibodies due to its interaction with the cellular receptor, C-C motif chemokine receptor 2 (CCR2) (L. Zhang et al., 2023). Studies indicate that the presence of anti-Gn neutralizing antibodies is associated with protection against SFTSV infection in experimental animal models and correlates with better clinical outcomes in humans (Chung et al., 2022; J.-Y. Kim et al., 2023; Song et al., 2018; S. Zhang et al., 2025). Considering these findings, we evaluated the presence of neutralizing antibodies in the sera of immunized mice; however, no significant neutralization activity was detected at 18 weeks post-priming for any of the experimental groups (data not shown).

Together, these findings indicate that the BPL inactivated SFTSV vaccine elicits a robust humoral response, and this response persists after 18 weeks post-priming. Our results also indicate an unbalanced antigen focus response, with the NP nucleoprotein the main antigenic focus of the humoral response. On the other hand, the glycoprotein Gn focused antibodies were poorly induced and were strongly dependent on the adjuvanted formulation. Notably, although not always statistically significant, the Addavax and SDI emerge as the most promising formulations with the most favorable overall performance.

### 4. Humoral response after booster immunization

Three weeks after booster immunization, all vaccine formulations elicited detectable antibody responses against whole SFTSV and the nucleoprotein NP. In contrast, reactivity against the Gn head domain remained comparatively modest in most groups (Figure 2C). Similar to the priming, the antibody responses against the whole virus, the NP nucleoprotein, and Gn glycoprotein were highest in the Addavax and SDI-adjuvanted groups, although these differences did not always reach statistical significance. In contrast, the IMDQ-P-C group showed significantly lower reactivity than both the Addavax and SDI-adjuvanted groups (p < 0.05). Differences among adjuvant formulations were more pronounced for the Gn head domain. In particular, the SDI-adjuvanted formulation induced the highest Gn head-specific antibody response and was significantly higher than all other groups except Addavax (p < 0.05; Figure 2C).

Comparison of antibody responses before and after booster immunization showed a significant increase in reactivity against whole SFTSV and nucleoprotein NP, whereas the increase in Gn-specific reactivity was limited and did not reach statistical significance (Supplementary Figure 2). In line with the enhanced antibody response directed against the Gn glycoprotein observed in the SDI and Addavax groups, serum neutralizing activity was detected exclusively in these groups (Figure 2D). The neutralizing antibody titers were higher in the SDI group than in the Addavax group (NC values of 81.6 and 33.1, respectively, expressed as reciprocal serum dilutions).

These findings are in line with the obtained 18 weeks after prime, indicating that i) NP represents an immunodominant antigenic hotspot and ii) among the formulations evaluated, SDI and Addavax were the most effective at inducing functional antibody responses after booster immunization. In addition, both formulations induce antibodies with broader reactivity, which include the Gn glycoprotein, and neutralizing activity under the conditions tested.

### 4. IgG isotype and Th1/Th2 skewed immune response

The analysis of IgG isotype profiles is a valuable tool for the qualitative characterization of immune responses induced by vaccination, providing insight into the immunological orientation elicited by different vaccine formulations or adjuvants. In murine models, a predominance of IgG1 is typically associated with a Th2 helper response, characterized by enhanced antibody production and strong humoral immunity. Elevated IgG2a levels associate with a Th1-skewed response, correlated with cellular immunity and the clearance of intracellular pathogens. We conducted IgG isotype profiling and observed marked differences in isotype distributions induced by the different formulations (Figure 2E). Alhydrogel and Addavax exhibited a pronounced and moderate skewing towards the IgG1 isotype, respectively, consistent with a Th2-biased response. SDI induced a balanced response, with a slight predominance of IgG2a, indicative of a Th1-skewed response. IMDQ-P-C alone and its combination with SDI elicited a strong bias toward IgG2a, accompanied by only modest IgG1 responses, suggesting a pronounced Th1 orientation, as previously described (Jangra et al., 2022).

### 5. Antigen-specific cellular memory response

A hallmark of immunological memory is the ability of primed T cells to undergo recall responses upon antigen reencounter. To determine whether the BPL-inactivated vaccine elicited persistent antigen-specific cellular recall, we performed an *ex vivo* antigen recall assay. At seven weeks post-boost, all animals were euthanized, and their spleens were harvested for further evaluation. The splenocytes were then isolated and stimulated *ex vivo* using the complete BPL-inactivated virus (SFTSV), the recombinant NP nucleoprotein, or the head domain of the Gn glycoprotein to assess the capacity of primed T cells to produce effector cytokines. Cytokine concentrations in culture supernatants were quantified by multiplex immunoassay and expressed as the antigen-induced change in cytokine concentration relative to the matched unstimulated condition (Δ cytokine concentration).

To evaluate the overall antigen-specific recall response, delta cytokine concentrations for each antigen-cytokine combination were compared against zero using two-sided one-sample Wilcoxon signed-rank tests. Thirty independent comparisons resulting from the three antigens (SFTSV, NP, and Gn head) and 10 cytokines (IL-1b, IL-2, IL-4, IL-5, IL-6, IL-12p70, IL-13, IL-18, IFN-γ, and TNF-α) were evaluated P-values were adjusted using the Benjamini-Hochberg procedure to control the false discovery rate (Supplementary Table 1). NP stimulation induced the broadest global recall response, with significant positive changes in 9 of the 10 measured cytokines after multiple-testing adjustment (q < 0.05; Supplementary Table 1). Ranked by median antigen-induced response, the strongest increases were observed for IL-6, IL-18, IFN-γ, IL-5, TNF-α, and IL-12p70, indicating a marked bias to Th1- and pro-inflammatory-type response. Interestingly, IL-2 did not show a significant global increase following NP stimulation. The whole inactivated SFTSV also induced a broad but lower-magnitude global recall response, with significant positive changes hierarchically ordered in IL-2, IL-5, IL-18, IL-13, IL-4, IL-6, after multiple-testing adjustment (all q < 0.05; Supplementary Table 1). In contrast, Gn head stimulation induced a limited and heterogeneous response. Only IL-2 showed a significant positive change, whereas IL-4 and TNF-α showed significant negative changes relative to matched unstimulated cultures; all remaining cytokines did not differ significantly from zero after multiple-testing adjustment. Together, these results indicate that the inactivated SFTSV vaccine elicited persistent antigen-specific immune memory detectable seven weeks after booster immunization and highlight NP as a major target of this response.

Although most between-adjuvant comparisons were not statistically significant, the distribution of cytokine responses following whole-virus stimulation suggested formulation-specific recall patterns (Figure 3). Addavax showed the most evident Th2-associated recall profile, characterized by prominent median responses for IL-4, IL-5, and IL-13, together with a strong IL-2 response. Notably, IL-4 was markedly induced, and a statistically significant difference was observed following whole-virus stimulation (versus IMDQ-P-C and IMDQ-P-C plus SDI: adjusted p=0.01 and 0.02, respectively). This pattern was consistent with the strong humoral responses elicited by the Addavax-formulated vaccine after both priming and booster immunization (Figures 2B and 2C), as well as with the prominent IgG1 response observed after boosting (Figure 2E). Similarly, Alhydrogel showed a Th2-associated recall pattern, with particularly prominent IL-5 and IL-13 responses, but less pronounced IL-4 induction than Addavax.

**Figure 3.**
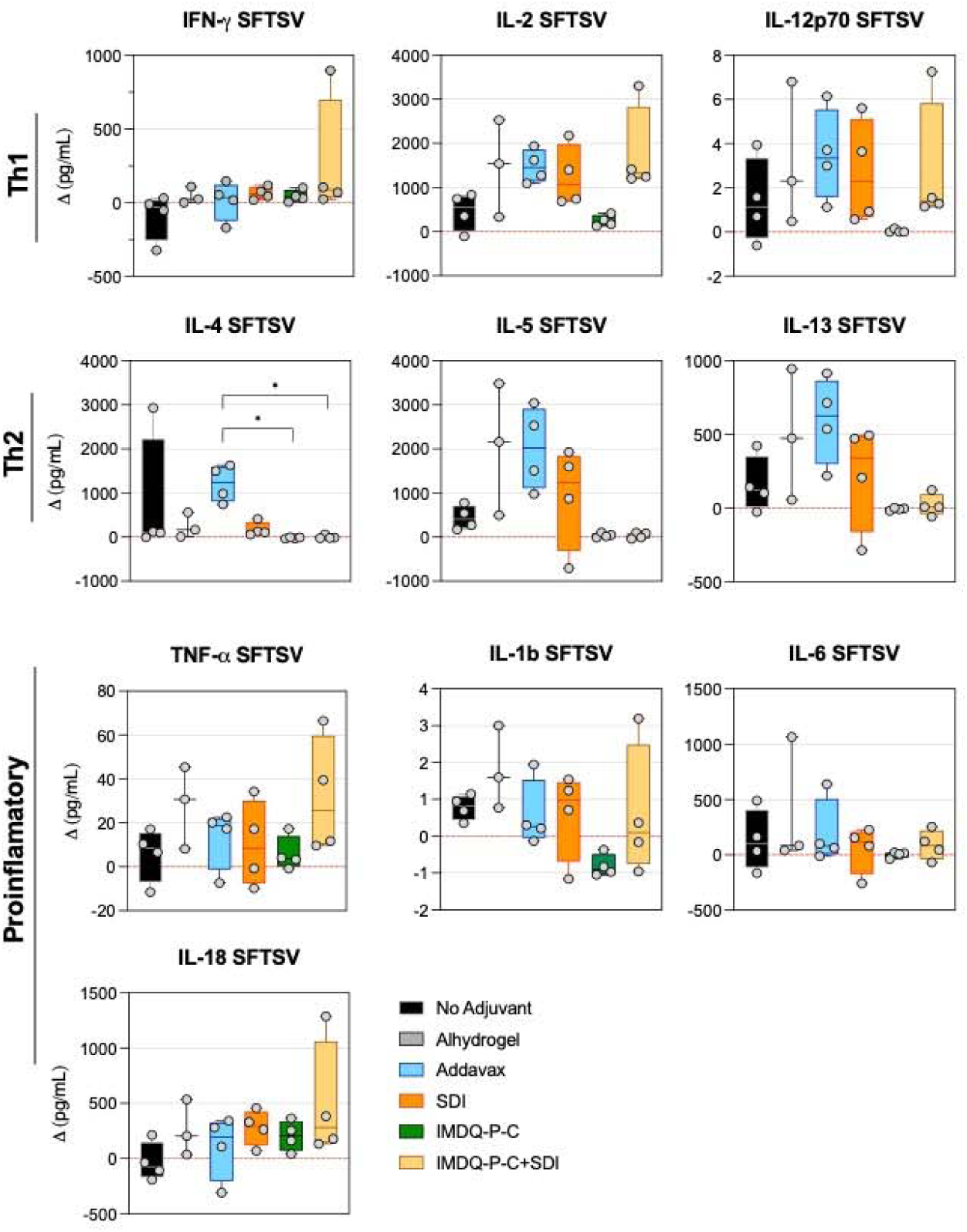
Cytokine expression following SFTSV whole antigen T cell recall. Splenocytes isolated from vaccinated mice were stimulated ex vivo with whole BPL-inactivated SFTSV. Cytokine concentrations in culture supernatants were quantified by multiplex immunoassay and expressed as antigen-induced changes relative to matched unstimulated cultures (Δ cytokine concentration = stimulated - unstimulated; picograms/mL). Individual points represent biological replicates (n = 4 per formulation, except Alhydrogel, n = 3 following exclusion of one technical failure). Cytokines are grouped as Th1-associated (top), Th2- associated (middle), and proinflammatory-associated (bottom). Box plots show the median and interquartile range; whiskers indicate the full data range. The dashed horizontal red line indicates Δ = 0, corresponding to no change relative to the matched unstimulated condition. Between- adjuvant comparisons were assessed separately for each cytokine using the Kruskal-Wallis test followed by Dunn’s multiple-comparisons test.

On the other hand, SDI displayed a more mixed whole-virus recall profile, combining IL-2 and Th2-associated cytokine responses with relatively higher IL-18 and modest IFN-γ induction. In contrast, the SDI plus IMDQ-P-C formulation showed an IL-2 and IL-18-associated profile with limited induction of Th2-associated cytokines. Although this group showed the highest median IFN-γ response following whole-virus stimulation, the absolute magnitude of IFN-γ induction remained modest (Figure 3).

Following *ex vivo* stimulation with recombinant NP, all vaccine formulations showed broad cytokine recall profiles characterized by concomitant Th1-associated, Th2-associated, and proinflammatory-associated responses (Figure 4). Although no between-adjuvant comparisons remained statistically significant after correction for multiple testing, the distribution of responses suggested formulation-specific patterns. IMDQ-P-C and the SDI plus IMDQ-P-C combination showed profiles relatively enriched in IFN-γ, IL-6, and IL-18, with concurrent increases in proinflammatory-associated cytokines, including IL-1β, TNF-α, and IL-12p70. These patterns were consistent with broad NP-induced cellular recall responses with relatively prominent Th1-associated and proinflammatory-associated features.

**Figure 4.**
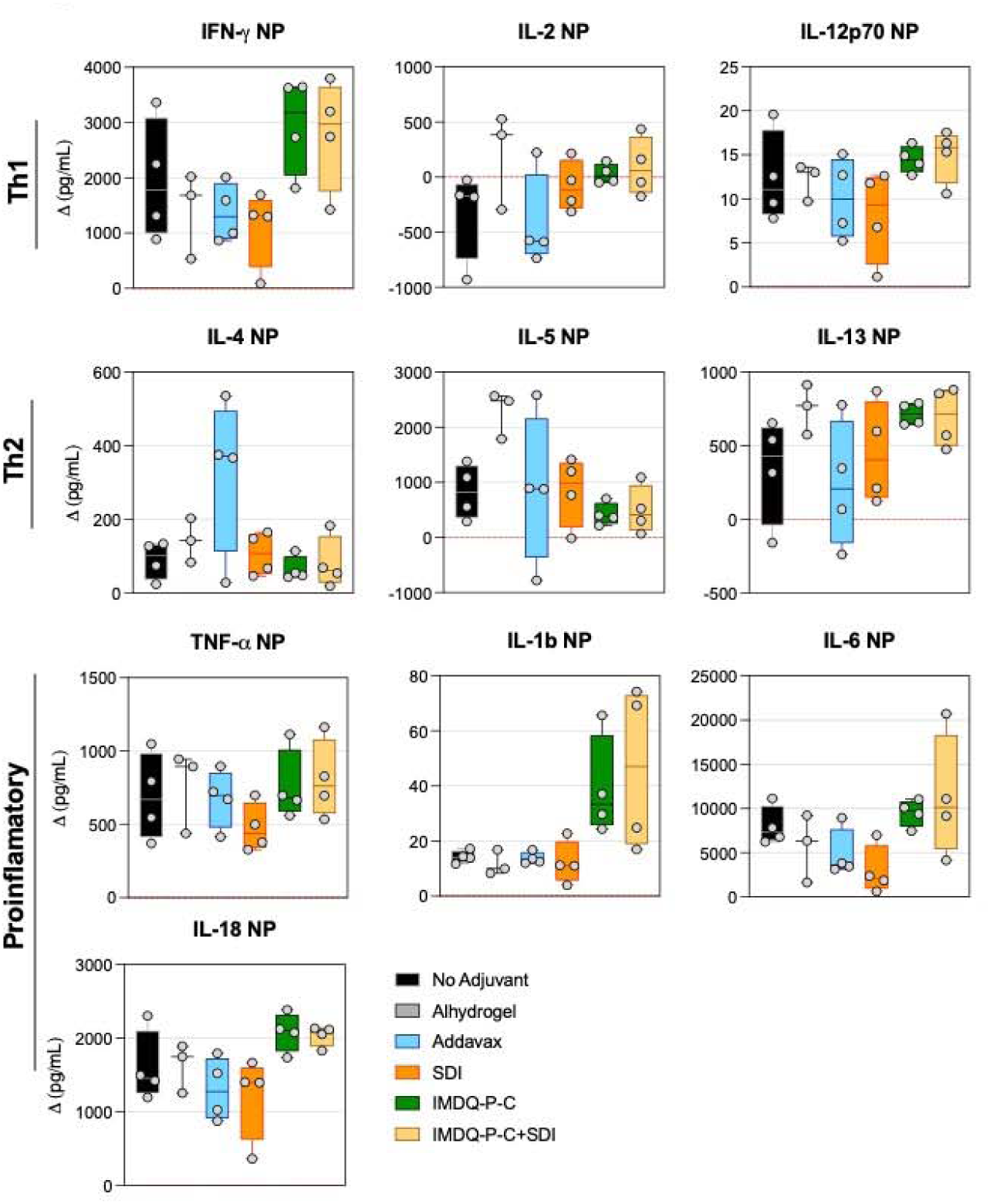
Cytokine expression following nucleoprotein NP T cell recall. Splenocytes isolated from vaccinated mice were stimulated ex vivo with recombinant SFTSV nucleoprotein (NP). Cytokine concentrations in culture supernatants were quantified by multiplex immunoassay and expressed as antigen-induced changes relative to matched unstimulated cultures (Δ cytokine concentration = stimulated − unstimulated; assay units). Individual points represent biological replicates (n = 4 per formulation, except Alhydrogel, n = 3 following exclusion of one technical failure). Cytokines are grouped as Th1-associated (top), Th2-associated (middle), and proinflammatory-associated (bottom). Box plots show the median and interquartile range; whiskers indicate the full data range. The dashed horizontal red line indicates Δ = 0, corresponding to no change relative to the matched unstimulated condition. Between-adjuvant comparisons were assessed separately for each cytokine using the Kruskal-Wallis test followed by Dunn’s multiple-comparisons test.

In contrast, Alhydrogel showed a broad mixed NP recall profile with a particularly prominent Th2-associated component, reflected by high IL-5 and IL-13 responses together with detectable IFN-γ, IL-6, IL-18, and TNF-α induction (Figure 4). Addavax also induced a mixed NP recall profile, characterized by comparatively prominent IL-4 responses in addition to measurable IFN- γ and proinflammatory-associated cytokine induction. SDI elicited an intermediate mixed profile, with concurrent IFN-γ, IL-5, IL-13, and IL-18 responses (Figure 4). Notably, substantial NP-induced cytokine responses were also observed in the non-adjuvanted vaccine group, supporting NP as a dominant recall antigen within this inactivated whole-virus vaccine platform.

In contrast to whole-virus and NP, *ex vivo* stimulation with the Gn head domain elicited low-magnitude and heterogeneous cytokine responses across vaccine formulations (Figure 5). Although selected groups showed modest increases in individual cytokines, including IL-2, IFN-γ, or IL-18, no coherent Th1-associated, Th2-associated, or proinflammatory-associated recall profile was evident. Addavax and SDI showed relatively higher IL-2 responses, whereas SDI showed modest IFN-γ and IL-18 responses; however, these patterns were not accompanied by coordinated induction of additional cytokines within a defined response profile. Consistent with the global analysis, Gn head stimulation did not elicit a broad antigen-specific recall response. Instead, several cytokines remained close to zero or showed negative delta values relative to matched unstimulated cultures, particularly for IL-4, IL-5, IL-6, and TNF-α in selected formulations (Figure 5). This limited and variable cellular recall pattern was consistent with the low Gn head-specific humoral responses observed after both priming and booster immunization (Figure 2 B and 2C). Overall, these findings indicate that the Gn head domain elicited limited antigen-specific cellular and humoral responses and did not reveal a clear formulation-specific cytokine signature.

**Figure 5.**
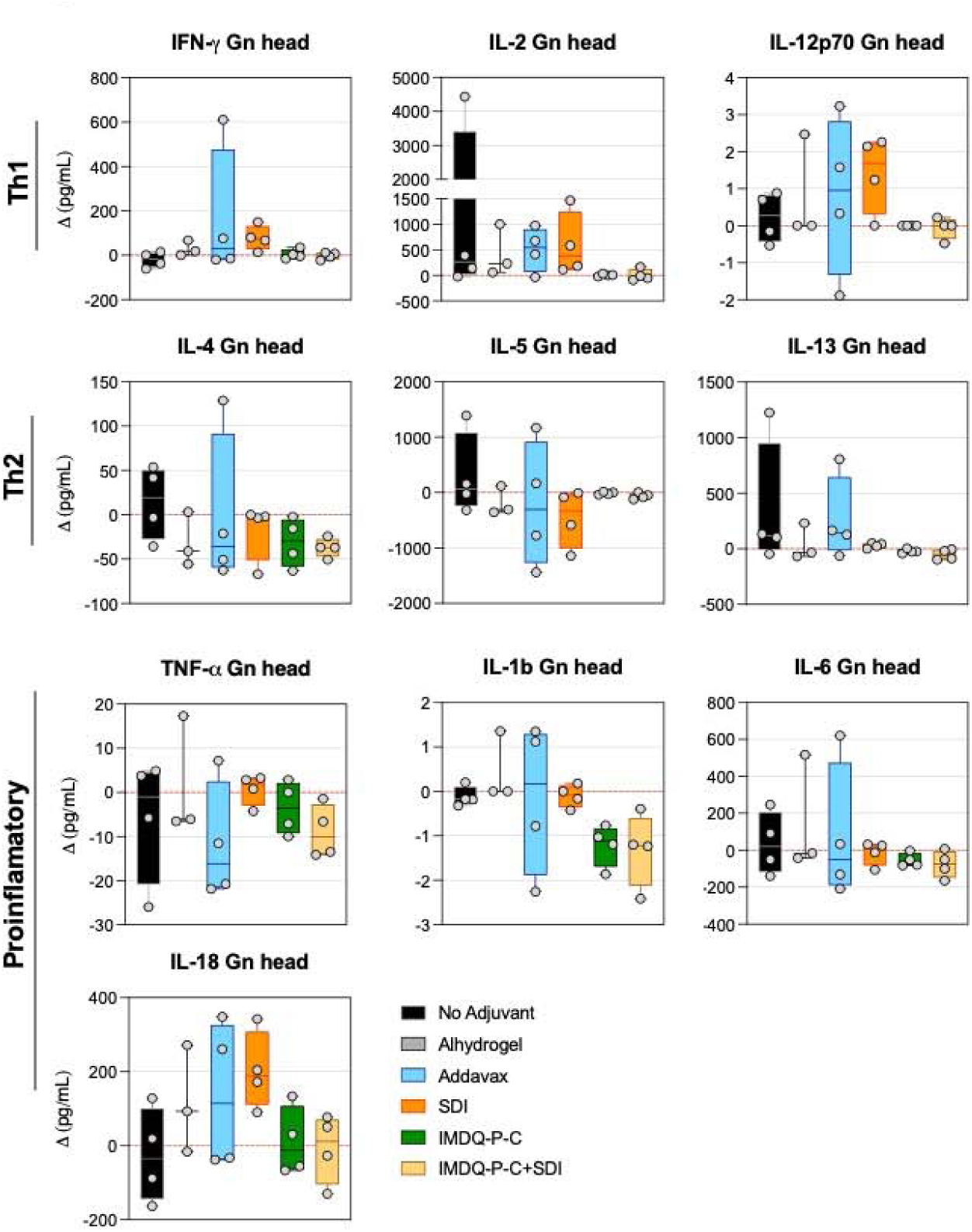
Cytokine expression following glycoprotein Gn T cell recall. Splenocytes from vaccinated mice were stimulated *ex vivo* with the recombinant head domain of the glycoprotein Gn. Cytokine concentrations in culture supernatants were quantified by multiplex immunoassay and expressed as antigen-induced changes relative to matched unstimulated cultures (Δ cytokine concentration = stimulated - unstimulated; assay units). Individual points represent biological replicates (n = 4 per formulation, except Alhydrogel, n = 3 following exclusion of one technical failure). Cytokines are grouped as Th1-associated (top), Th2-associated (middle), and proinflammatory-associated (bottom). Box plots show the median and interquartile range; whiskers indicate the full data range. The dashed horizontal red line indicates Δ = 0, corresponding to no change relative to the matched unstimulated condition. Between-adjuvant comparisons were assessed separately for each cytokine using the Kruskal-Wallis test followed by Dunn’s multiple-comparisons test.

### 6. Vaccine-Induced Humoral Protection *In Vivo*

To assess the protection offered by the humoral responses induced by different vaccine formulations, we conducted a passive serum transfer in type I interferon receptor-deficient mice (IFNAR-/-) followed by a lethal viral challenge. This mouse model was selected due to its inability to mount effective innate or adaptive immune responses, resulting in uncontrolled viral replication and lethality upon infection.

To validate our model, a pilot experiment was conducted in IFNAR-/- mice challenged with either 1 or 10 TCID of virus. Infected animals exhibited progressive weight loss beginning on day 3 (Supplementary Figure 3A). By day 6, mice were euthanized, and viral RNA levels were quantified in blood and spleen. High viral loads were detected in blood (geometric means of 2.84 × 10 and 2.1 × 10 genome copies per mL) and spleen (geometric means of 3.47 × 10 and 5.42 × 10 genome copies per gram of tissue) in the 1 and 10 TCID groups, respectively (Supplementary Figure 3B and 3C). Based on these results, an intermediate challenge dose of 5 TCID was selected for subsequent experiments.

The previously primed and boosted BALB/c mice (n=5 per group) were euthanized at seven weeks post boost, and total blood was obtained by intracardiac puncture (Figure 2A). All vaccinated mice showed high antibody titers against the whole virus (Figure 6A). Next, pools of sera from each group were prepared, and the anti-SFTSV endpoint titers of each pool were determined, showing concordance with the individual titers (Figure 6A).

**Figure 6.**
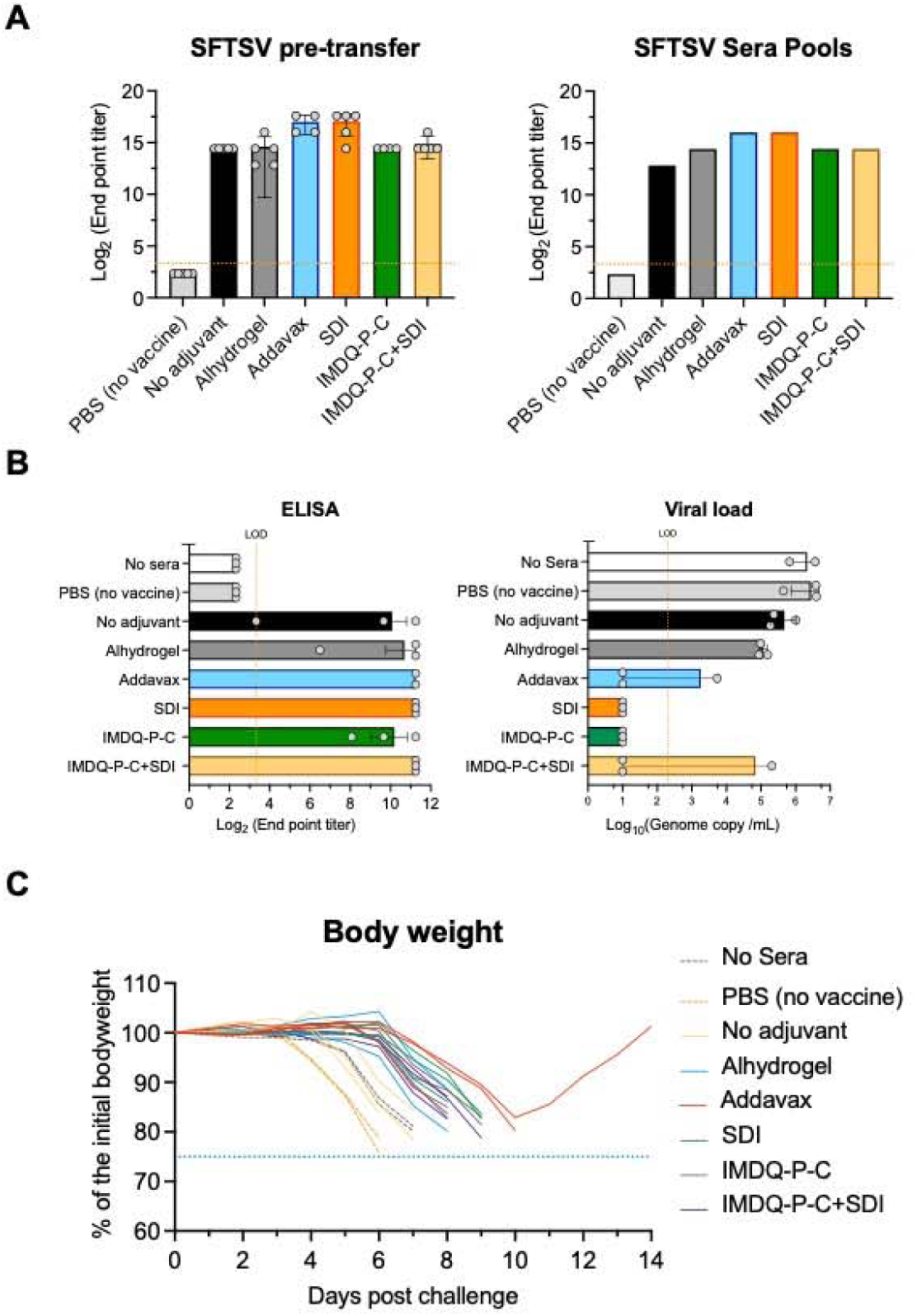
Passive transfer. **A.** End point titers of anti-SFTSV antibodies obtained from terminal intracardiac bleeding, determined by ELISA (left graph). Bars represent the median and interquartile range (IQR) of endpoint reciprocal dilution; individual dots are also shown. The dotted line represents the limit of detection (LOD). Antibody endpoint titers from sera pools from immunized mice (right graph). The bars represent the absolute value of anti-SFTSV antibody titer of each condition (adjuvanted formulation). The dotted line represents the limit of detection. **B.** Comparative analysis of antibody titers and viral load at day 5 post-challenge. Groups of IFNAR-/- mice (n=3) were passively immunized with pooled serum derived from immunized donor mice and subcutaneously challenged with 5 TCID□□ of SFTSV. Blood samples were collected on day 5 post-infection. Anti-SFTSV antibody titers and viral RNA genome copies/mL were quantified by ELISA and RT-qPCR, respectively. Data are presented as horizontal bars indicating the mean and standard deviation (SD). The dotted line marks the assay’s limit of detection. **C.** Following infection, mice were monitored daily for general health and body weight. The graph displays individual weight trajectories, with each line color-coded to represent the respective adjuvanted or control group. The blue dotted line indicates the humane endpoint threshold.

IFNAR-/- mice (n=6 per group; 3 males and 3 females) received 250 µL of pooled serum via intraperitoneal injection (day 0). Two days later, mice were challenged subcutaneously with 5 TCID of virus to simulate a natural infection via a tick bite.

At day 5, blood samples were collected to confirm the presence of transferred antibodies and to quantify viral loads. Antibody titers in recipient mice correlated with those of the transferred sera, with higher reactivity observed in mice that received sera from the Addavax and SDI groups (Figure 6B). Importantly, these groups also showed lower viral loads, exhibiting an inverse relationship between antibody titers and viral replication (Figure 6B).

Dairy monitoring revealed the onset of weight loss between days 3 and 7 post-challenge, with the timing of disease progression and mortality varying depending on the serum donor group (Figure Supplementary 4). Mice in the no vaccine donor group and the no-serum (PBS) control groups were the first to exhibit signs of illness, beginning as early as day 3 to 4, with complete mortality by day 7 and 8, respectively, indicative of a total lack of protection (Figure 6C and Supplementary Figure 4).

Among the groups that received serum from BPL-inactivated vaccine recipients, the non-adjuvanted donor group provided the lowest protection, with the onset of clinical symptoms occurring at days 3-4, and 100% mortality was observed by day 7 (mean survival end point=6.67 days, Supplementary Figure 4). In contrast, the SDI, IMDQ-P-C, and IMDQ-P-C + SDI groups conferred a moderate degree of protection, with an onset of clinical symptoms at day 6, delaying mortality until day 9 (mean survival endpoint: 8.667, 8.67, and 8.33 days, respectively). Notably, the Addavax receptor group demonstrated the most protective effect, showing the clinical onset at 6-7 days, and more importantly, it was the only condition that conferred measurable survival, with one mouse remaining alive at day 14 (Figure 6C and Supplementary Figure 4, mean survival endpoint >10.667 days). The delayed onset of clinical signs, the reduced viral loads, and prolonged survival, especially those receiving sera from adjuvanted formulations, highlight the crucial role of adjuvant choice in determining the quality and effectiveness of the humoral response.

### Cross-reactivity of the vaccine humoral response

Finally, we evaluated the cross-reactivity of vaccine-induced sera against other bunyaviruses, including a homologous SFTSV strain (HB29) and a heterologous virus from a different genus (Heartland virus, strain MO4). Microneutralization assays were performed using sera from all adjuvanted vaccine groups against Heartland virus; however, none of the samples exhibited neutralizing activity (Figure 7A). This outcome was anticipated, as Heartland virus and SFTSV belong to different genera within the *Phenuiviridae* family, and their structural antigens are highly divergent.

**Figure 7.**
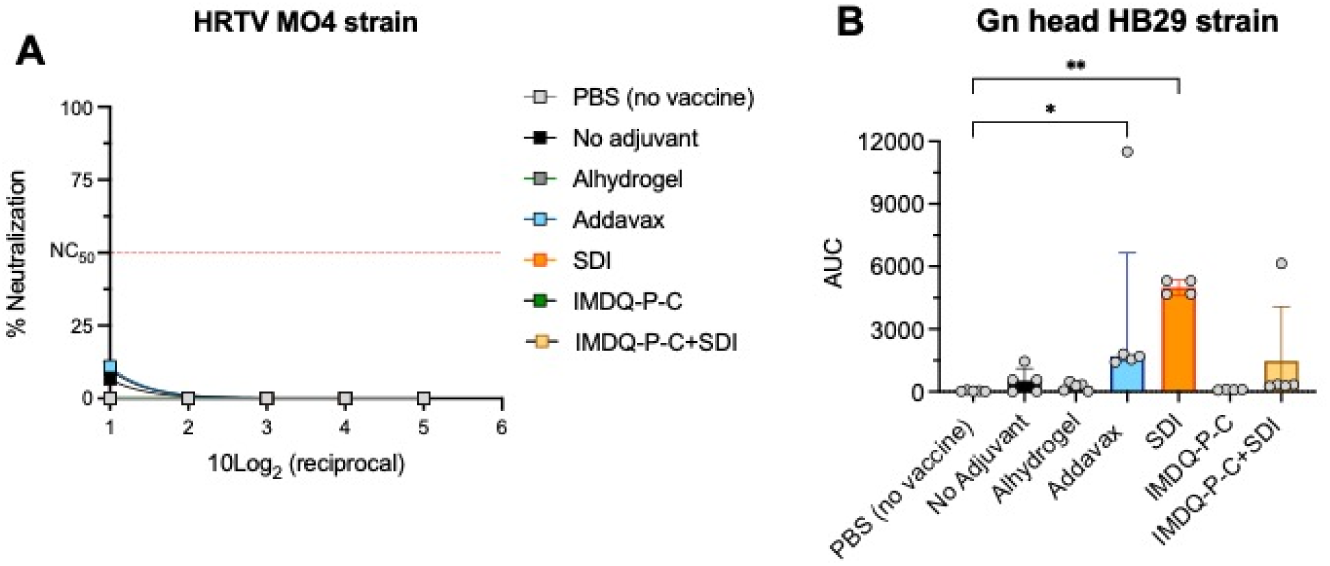
Cross-reactivity of the humoral response in vaccinated mice. **A.** Neutralization assay was performed using 100 TCID□□ of Heartland virus strain MO4 incubated with two-fold serial dilutions of sera obtained from vaccinated mice, collected 7 weeks post-boost. After 4 days of incubation, infection was assessed by a plaque-forming assay, which visualized cytopathic effects through crystal violet staining after fixation with 7% paraformaldehyde. Samples were analyzed in duplicate from five biological replicas. **B.** Mice sera collected 7 weeks post-boost were tested for reactivity against the recombinant Gn glycoprotein of the SFTSV HB29 strain. The mean and SD of the area under the curve (AUC) for each vaccinated or control group are shown. The dotted line indicates the limit of detection (LOD).

Remarkably, we detected serum cross-reactivity against the recombinant Gn glycoprotein of SFTSV strain HB29 in sera from mice immunized with SDI and Addavax-adjuvanted formulations in ELISA (Figure 7B). Although neutralization assays with HB29 could not be conducted due to limited serum availability, the observed serological reactivity is noteworthy.

## Discussion

The recognition of SFTSV as a significant public health threat has prompted various research efforts to develop a vaccine, adopt multiple strategies, and demonstrate the feasibility of these approaches. Reports indicate that live-attenuated, DNA, mRNA, protein subunit, nanoparticle, and viral vector vaccines all offer distinct levels of protective immunity (Lu et al., 2024; S. Kim et al., 2024; K.-M. Yu et al., 2019a; Kwak et al., 2019; Zhao et al., 2020; D. Kim et al., 2023; Jeong et al., 2025; K.-M. Yu et al., 2019b).

This study outlines a practical approach for developing a BPL-inactivated vaccine against SFTSV, underscoring the importance of adjuvant selection in shaping immune outcomes. Importantly, the production of high viral titers under serum-free conditions, combined with PEG-based precipitation and BPL inactivation, offers a practical and scalable strategy suitable for future large-scale vaccine manufacturing.

Consistent with previous reports, treatment with 0.05% BPL was sufficient to completely inactivate our viral strain, as confirmed by viability assays showing no residual infectivity (Gupta et al., 2021; Roberts et al., 2010). Furthermore, the robust humoral response observed after both the prime and boost immunizations suggests that the structural and antigenic integrity of the virus was preserved.

A distinctive feature of this study was the extended 21-week interval between priming and booster immunization. Although prolonged prime-boost intervals have not been systematically evaluated for SFTSV vaccine candidates, studies in other vaccine platforms have shown that extending the interval between doses can influence immune maturation, the durability and quality of humoral and cellular responses, inflammatory programs, and recall capacity (Ahmed et al., 2025; Amini et al., 2025; Murray et al., 2025; Nicolas et al., 2023; Nielsen et al., 2023; Pettini et al., 2021). In the present study, the extended interval provided an exploratory framework to assess the persistence of vaccine-induced immunity before boosting and the capacity to mount antigen-specific recall responses several weeks after the booster dose. The robust post-boost antibody responses, together with the broad specific cellular recall observed seven weeks after boosting, indicate that immunological responsiveness was retained across this prolonged interval.

Further evaluation of humoral responses showed that adjuvanted formulations significantly increased antibody titers. Addavax and SDI elicited strong antibody responses, especially against NP and the head domain of the Gn glycoprotein, including functional antibodies with neutralizing activity. In contrast, IMDQ-P-C and its combination with SDI induced comparatively weaker humoral responses. However, as indicated by IgG isotyping, these last two formulations promoted a Th1-skewed profile, characterized by elevated IgG2a levels. This polarization, although associated with lower antibody magnitude, may be particularly beneficial for supporting cellular immunity and promoting viral clearance. Meanwhile, more balanced isotype profiles induced by SDI alone or Addavax could offer advantages in controlling immunopathology, especially in infections such as SFTSV, where excessive inflammation contributes to disease severity. These findings underscore the importance of aligning adjuvant-induced immune profiles with both the protective and regulatory needs of the target disease (Bretscher, 2019; Clerici & Shearer, 1993; Gil-Etayo et al., 2021).

Similarly, the antigen recall cytokine response was modulated by the different adjuvants, and the distribution of cytokine responses suggested formulation-specific patterns. For example, after whole-inactivated virus stimulus, Addavax showed the most evident type 2-associated profile, with prominent IL-4, IL-5, and IL-13 responses together with strong IL-2 induction. This pattern was consistent with the strong humoral responses induced by the Addavax-formulated vaccine after priming and booster immunization, as well as with the prominent IgG1 response observed after boosting. Alhydrogel showed a broadly similar pattern, with prominent IL-5 and IL-13 responses but less marked IL-4 induction than Addavax. SDI showed a more mixed whole-virus recall profile, combining IL-2 and type 2-associated cytokine responses with relatively higher IL-18 and modest IFN-γ induction. In contrast, the SDI plus IMDQ-P-C formulation showed an IL-2- and IL-18-associated profile with limited induction of IL-4, IL-5, and IL-13. Although this group showed the highest median IFN-γ response after whole-virus stimulation, the absolute magnitude of IFN-γ induction remained modest.

A remarkable observation from our study was the differential modulation of the immune responses by the different viral structural components. Unfortunately, we were not able to detect a humoral response against the Gc glycoprotein, potentially due to the lack of structural fidelity of our recombinant glycoprotein. It is essential to mention the relevance of confirming this factor in the design of subunit vaccines or the implementation of serological tests, i.e., ELISA. In our hands, we observed Gc reactivity only in the cellular context, suggesting that, like other bunyaviruses, intermolecular interactions with Gn and the lipidic membrane have a significant influence on Gc stabilization. This phenomenon is particularly relevant to vaccine design, as novel mRNA vaccines targeting the Gc glycoprotein may elicit antibodies against improperly folded antigens. The implications of this process warrant further investigation. (Du et al., 2023; Plegge et al., 2016; Z. Sun et al., 2023; X. Zhang et al., 2025).

In contrast, the Gn glycoprotein does not present antigenic fidelity issues, potentially because its head domain is structurally independent of such interactions and displays a stable surface interaction in the soluble phase (Du et al., 2023; Z. Sun et al., 2023). Our analyses suggest that the Gn glycoprotein exhibits limited immunogenicity in the context of the inactivated virus vaccine. This may be partly attributed to the presence of two surface-exposed glycosylation motifs that can mask antigenic epitopes and modulate immune recognition (Tate, 1938).

Consistent with this, our ELISA results showed only modest antibody reactivity to Gn in vaccinated mice. Similarly, the cytokine response to Gn *ex vivo* stimulation was uniformly low across all vaccine formulations.

A central finding of this study was the dominance of NP as a target of vaccine-induced immune memory. Among the three recall antigens evaluated, NP elicited the broadest and highest-magnitude cytokine response, exceeding that induced by whole BPL-inactivated SFTSV and contrasting sharply with the limited response observed after Gn head stimulation. This pattern was consistent with the robust NP-specific antibody responses detected after both priming and booster immunization, indicating that NP was preferentially recognized by both the humoral and cellular arms of the vaccine-induced response. Although we did not directly assess the neutralizing capacity of NP-specific antibodies, their high abundance following priming, combined with the absence of neutralizing activity in the total serum, suggests that these antibodies lack neutralizing function in our model. These findings are in line with previous studies in which DNA vaccination targeting N elicited strong T cell responses and non-neutralizing antibodies, although protection remained partial (Kang et al., 2020; Kwak et al., 2019).

Although the immunodominance of the nucleoprotein remains unexplained, it is plausible that, as the most abundant structural component of the virion, its intrinsic immunogenicity is further amplified by its capacity to form complexes with viral RNA. In the context of a β-propiolactone (BPL)-inactivated whole-virus vaccine, residual RNA fragments bound to the N protein may remain structurally preserved and be recognized by cytosolic pattern recognition receptors such as the retinoic acid-like receptors RIG-I and mda5. Short 5’-triphosphorylated viral RNA molecules complexed with the N protein could be recognized by the RIG-I helicase receptor in the cytoplasm of host antigen-presenting cells. Upon recognition by RIG-I and through MAVS activation, downstream signaling pathways are triggered, including the activation of IRF3/7 and NF-κB, which results in the transcriptional induction of type I interferons and pro-inflammatory cytokines. This antiviral response enhances the maturation of dendritic cells and the priming of CD8 cytotoxic T lymphocytes through MHC class I-restricted antigen presentation, thereby supporting robust cellular immunity (Liu & Gack, 2020).

In contrast to NP, Gn induced weak antibody responses, and its immunogenicity was strongly formulation dependent. SDI and, to a lesser extent, Addavax were the only formulations that significantly induced Gn-specific antibodies after both priming and booster immunization.

Concomitantly, sera obtained from the SDI- and Addavax-formulated vaccine groups were the only ones that exhibited neutralizing activity.

Because the Gn glycoprotein has previously been described as a major target of neutralizing antibodies, these findings highlight the critical role of adjuvants in directing the antigenic focus of vaccine-induced humoral immunity. Whereas NP dominated the overall immunological response to the inactivated vaccine, SDI and Addavax demonstrated the capacity to broaden antibody recognition beyond this immunodominant target, promoting Gn-specific antibody responses and neutralizing activity. Collectively, these observations indicate that effective SFTSV vaccine design may require the complementary targeting of NP to support robust cellular immunity and Gn to promote neutralizing antibody responses.

Functional protection studies using passive serum transfer in IFNAR-/- mice further supported the role of antibody-mediated immunity. Sera from animals immunized with Addavax- and SDI-adjuvanted vaccines delayed disease onset, reduced viral loads, and conferred partial protection against lethal challenge. Notably, all mice in the experimental groups succumbed to infection except for a single recipient of Addavax-derived immune serum, who showed transient illness but ultimately recovered. This outcome highlights both the functional relevance of humoral responses and the limitations of the IFNAR-/- model, which, due to its lack of innate immune signaling, is highly susceptible to infection and sensitive to the magnitude of the initial viral inoculum. Despite these constraints, partial protection was evidenced by undetectable viral loads at day 5 post-challenge, delayed symptom onset (e.g., weight loss), and prolonged survival in recipients of adjuvanted sera compared to non-adjuvanted and non-vaccinated controls. These findings underscore the importance of antibody-mediated mechanisms and highlight Addavax as a promising adjuvant for future SFTSV vaccine development.

One limitation of this study is the lack of infection data for female recipients in the passive transfer and challenge experiments. However, prior data indicate that male and female IFNAR-/- mice exhibit comparable infection dynamics, with similar viral replication rates and clinical trajectories, characterized by initial weight loss around day 4 and mortality between days 6 and 8 post-challenge (Supplementary Figure 5). This suggests that the findings observed in males are likely to be broadly representative.

Of particular interest was the detection of cross-reactivity against the recombinant Gn glycoprotein of SFTSV strain HB29 in sera from mice immunized with SDI and Addavax- adjuvanted formulations. HB29 and the vaccine strain YL1 used for vaccination belong to different genotypes (D and F, respectively), which are associated with distinct clinical outcomes, HB29 being linked to higher mortality rates (Dai et al., 2022). Structural modeling indicates that six amino acid differences between their Gn glycoproteins cluster near the hypothetical receptor-binding site in subdomains I and III (Supplementary Figure 6). While two apical N-glycosylations mask the subdomain I, subdomain III is sterically free and is considered the hot spot for neutralizing antibodies. Importantly, YL1 and HB29 share a high similarity in subdomain II, which is less exposed on the capsid and is a plausible focus for cross-reactive antibodies. Our observations show that the adjuvants SDI and Addavax not only improved the general reactivity to SFTSV but also were able to elicit cross-reactive antibodies, although their antigenic targets remain to be determined. The inactivated-virus vaccines represent one of the oldest and most broadly applied strategies in vaccinology, with variable outcomes depending on the pathogen. Some, such as the inactivated poliomyelitis vaccine, have achieved extraordinary success, contributing to the control and near-eradication of the disease (Bandyopadhyay et al., 2024). Others have shown only modest or inconsistent efficacy (Q. Wu et al., 2023), and, in some cases, such as with early respiratory syncytial virus vaccines, have even exacerbated disease upon exposure to the pathogen, leading to severe outcomes and fatalities (Castilow et al., 2007). These contrasts highlight the need to improve the immunogenicity and protective efficacy of inactivated vaccines, thereby considering host immune response skewing using appropriate adjuvants. A promising approach involves the use of rationally selected adjuvants that activate innate immune sensors, such as TLR or RIG-I receptors, to boost both humoral and cellular immunity. In this study, we employed two such adjuvants, SDI and IMDQ-P-C, designed to target RIG-I and TLR7/8 pathways, respectively. Our findings demonstrate that these adjuvants markedly enhanced the immunological profile induced by the inactivated SFTSV vaccine.

In conclusion, this study establishes a comprehensive and rigorous experimental platform for the evaluation of SFTSV vaccine candidates, underscoring the critical interplay between antigen selection and adjuvant pairing in shaping effective immune responses. The synergy observed between Th1-skewing adjuvants, such as IMDQ-P-C, and adaptable immunomodulators like SDI provides a rational framework for modulating both humoral and cellular arms of immunity.

Notably, the consistent immunodominance and functional relevance of NP-specific responses support its inclusion as a central component in next-generation vaccine strategies. In parallel, this work demonstrates the effectiveness of Addavax in enhancing functional humoral immunity, reinforcing its value as a clinically validated adjuvant. Together, these findings offer critical conceptual and translational insights for the rational design and continued refinement of safe, efficacious, and broadly protective vaccines against this emerging tick-borne pathogen.

## Acknowledgements

The Conventional Biocontainment Facility (CBF) is a NIHBSL3/BSL3 facility that is part of the BSL-3 Biocontainment CoRE. This core is supported by subsidies from the ISMMS Dean’s Office and by investigator support through a cost recovery mechanism. Research reported in this publication was supported by the National Institute of Allergy and Infectious Diseases of the National Institutes of Health under Award Number G20AI174733 (R.A. Albrecht). The work in this study was funded by NIAID contract 75N93019C00046 to A.G-S.. V.Y. and L.A.C. were supported by NIH training grant T32 AI007647 and E.B. by NIH training grant T32 AI078892.

---

**Supplementary Figure 1.**
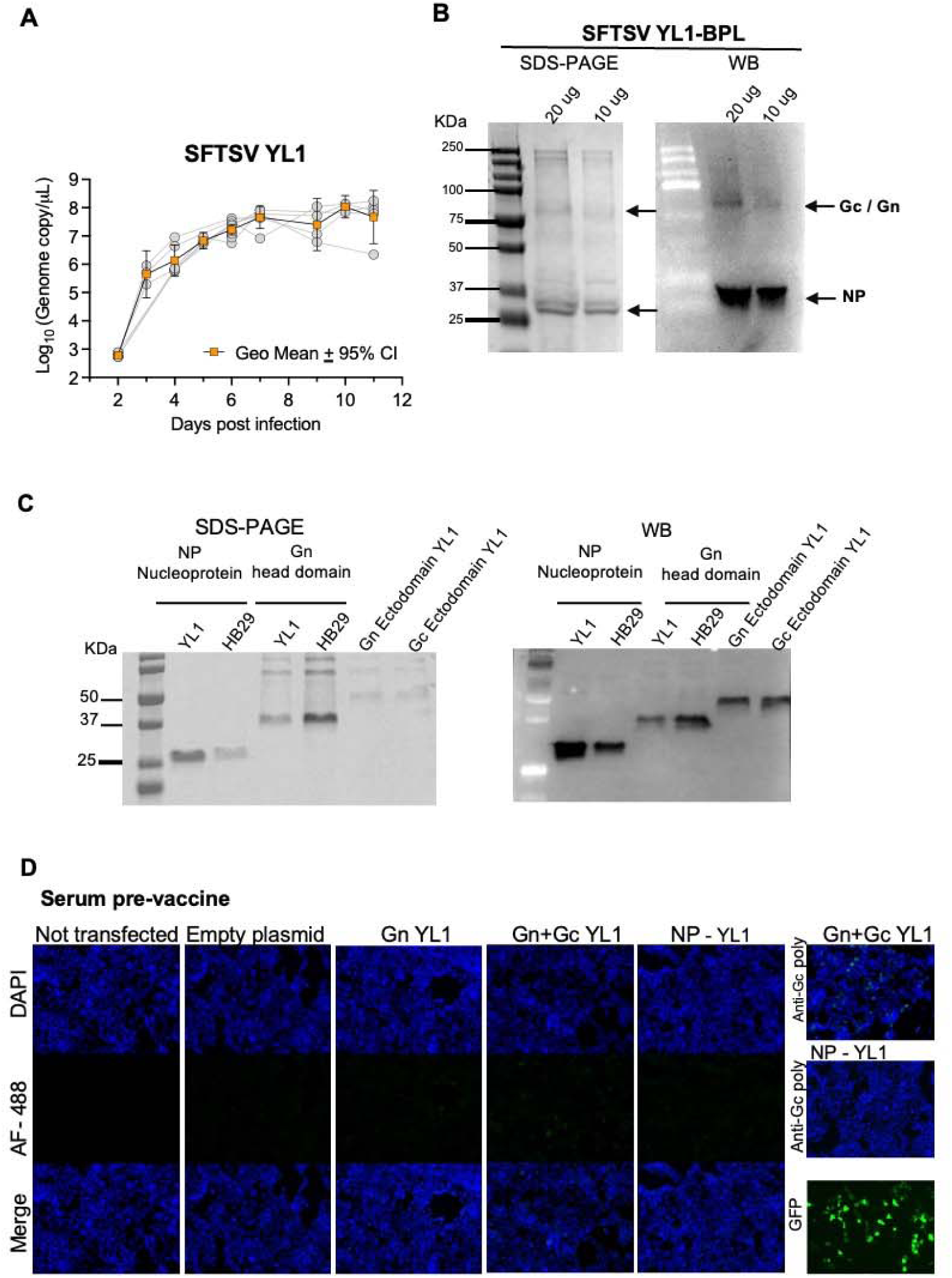
**A. Viral replication in serum-free medium.** T175 flasks were seeded with VERO E6 cells to obtain 70-80% confluence on the day of infection. The infection proceeds at a multiplicity of infection (MOI) of 0.01, and the culture medium OptiPRO SFM was adjusted to 30 mL. Samples of the culture supernatant were taken daily, and the SFTSV viral load (genome copies/mL) was determined by RTqPCR. The geometric mean and 95% confidence interval (CI) are shown. **B. Recombinant protein production.** A plasmid encoding the full open reading frame (ORF) of the nucleoprotein (NP) was expressed in E. coli, and the complete ectodomain and the head domain of the Gn glycoprotein, the complete ectodomain of glycoprotein Gc, in Expi293F mammalian cells. Proteins were purified using gravity-flow IMAC (Qiagen). Western blot analysis using anti-His tag antibodies showed the protein expression after purification. **C. Vaccine preparation.** SFTSV strain Y1 was propagated in five T175 as showed in A. After 7 days of incubation, viral supernatants were clarified and concentrated by precipitation with polyethylene glycol (PEG; Abcam, Fisher Scientific). Inactivation of the virus was performed using 0.05% β-propiolactone (BPL), and the absence of residual infectivity was confirmed by TCID□□ assay. The image displays the final vaccine preparation analyzed by denaturing SDS-PAGE. **D. Complete open reading frame (ORF) Glycoprotein Gc expression in HEK293 cells.** Gc. Panel shows the negative control of Figure 1B. Mouse serum from a pre-vaccinated mouse was checked for reactivity to Gc glycoprotein. As in 1B, the glycoprotein Gn and nucleoprotein NP (both complete ORF) were also expressed as controls. A commercial anti-Gc antibody was used as a reactivity control, and a GFP coding expression plasmid was used to check the transfection performance. Empty plasmid and non-transfected controls are also shown.

**Supplementary Figure 2.**
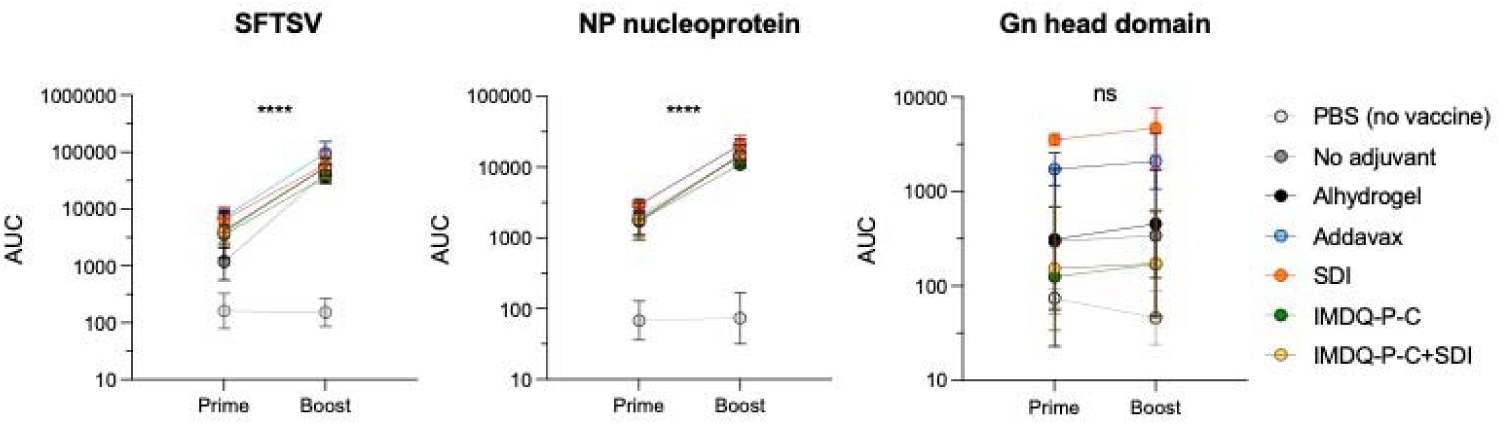
Prime vs boost humoral response comparison. The mean and standard deviation (AUC) values of the mice sera reactivity after prime and boost are displayed, and differences were assessed using one-way ANOVA, followed by Tukey’s multiple comparisons test for paired data. Statistical significance was defined as P < 0.05.

**Supplementary Figure 3.**
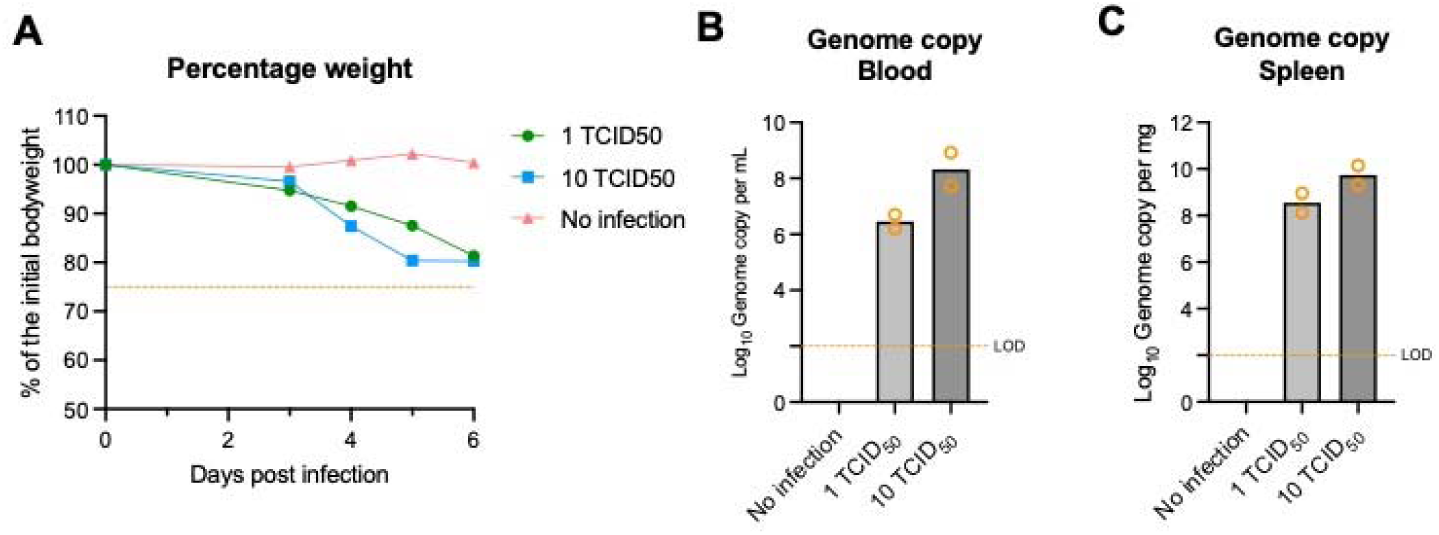
A. Mouse infection pilot experiments. Percent weight loss of mice infected subcutaneously with either 1 or 10 TCID□□ of SFTSV, or mock-infected controls. Each line represents the change in body weight over time relative to the initial weight (expressed as a percentage). Data points represent the mean of two biological replicates. The dotted line indicates the humane endpoint threshold, which is 75% of the initial body weight. **B.** Viral load in blood on day 5 post-infection. Blood samples were collected via retro-orbital puncture for quantification of viral loads (genome copies/mL) by RT-qPCR. The graph shows geometric mean values from two biological replicates; individual values are plotted. **C.** Viral load in spleen tissue on day 5 post-infection. Following euthanasia, spleens were harvested and weighed, and the viral loads were determined by RT-qPCR and expressed in genome copies per mg of tissue. Calculations and graphs were generated using GraphPad Prism.

**Supplementary Figure 4.**
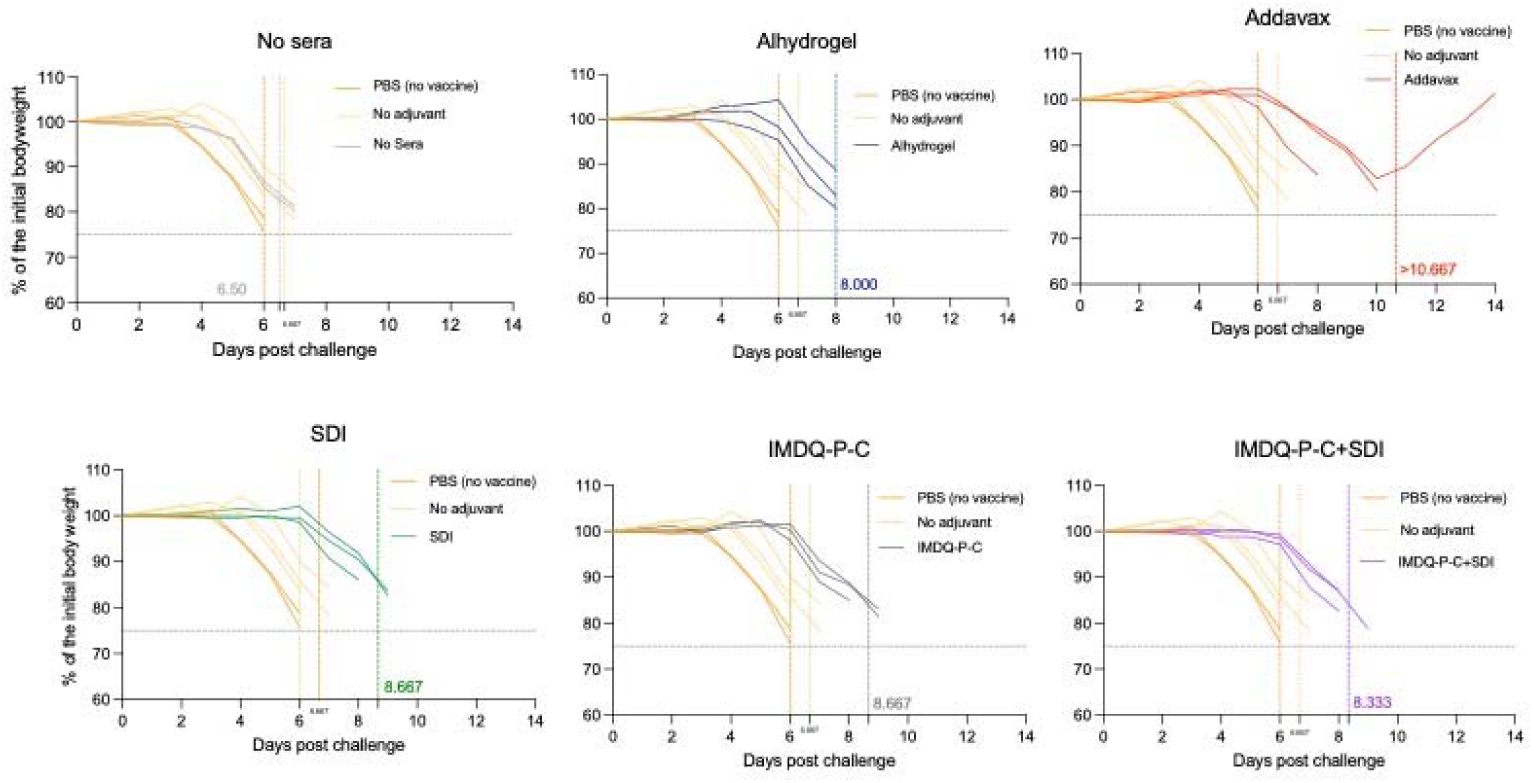
Body weight curves by donor vaccine formulation. After the infection, mice were monitored daily for general health and body weight. The graph displays individual weight trajectories, with each line color-coded to represent the respective adjuvanted or control group. The vertical line shown in the corresponding group color indicates the mean survival endpoint, defined as the mean final day on which animals in that group remained alive. The numerical value displayed at the base of each vertical line denotes the corresponding mean survival day. The horizontal dotted gray line indicates the humane endpoint threshold.

**Supplementary Figure 5.**
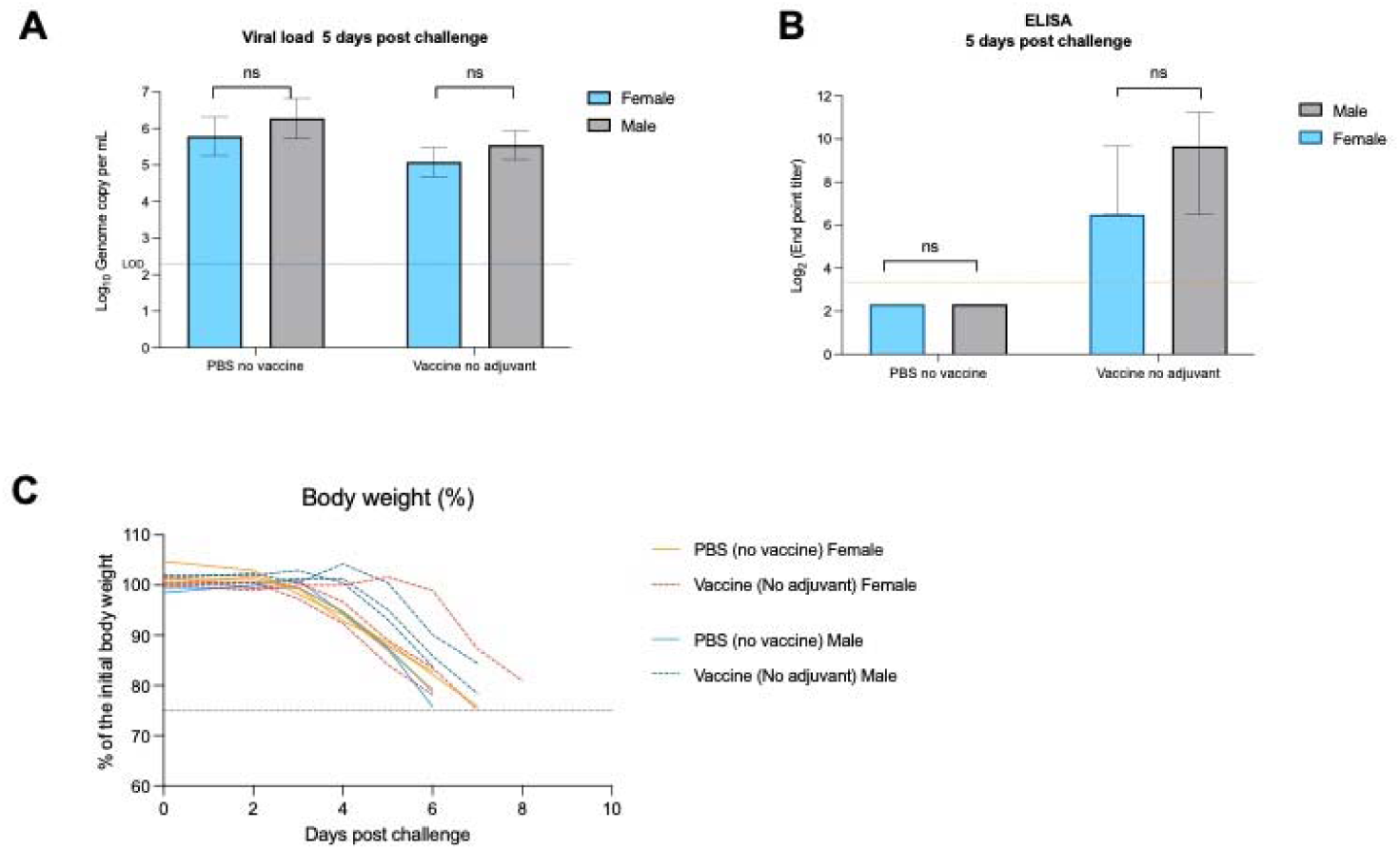
Comparison of the SFTSV infection course between male and female mice. **A.** Groups of three male and three female IFNAR-/- mice were infected subcutaneously with 5 TCID_50_ of SFTSV. Five days post-infection, viral loads in blood were determined by RT-qPCR. Bar graphs show geometric means ± standard deviation (SD). The dotted line represents the assay limit of detection. **B.** At 5 days post-infection, blood samples were collected, and subsequent ELISA using whole inactivated SFTSV as antigen was conducted on the serum samples. Endpoint titers are shown as the reciprocal of the last dilution. Bar graphs indicate the median and interquartile range (IQR). Parametric and non-parametric statistical tests were applied in panels A and B, respectively, to assess differences between groups. **C.** Following infection, mice were monitored daily for general health and body weight. The graph displays individual weight variations, with each line color-coded to represent the respective adjuvanted or control group. The dotted line indicates the humane endpoint threshold. For clarity, a nominal weight value of 60 was assigned to mice at the time of death. All animals succumbed to infection before reaching the humane endpoint.

**Supplementary Figure 6.**
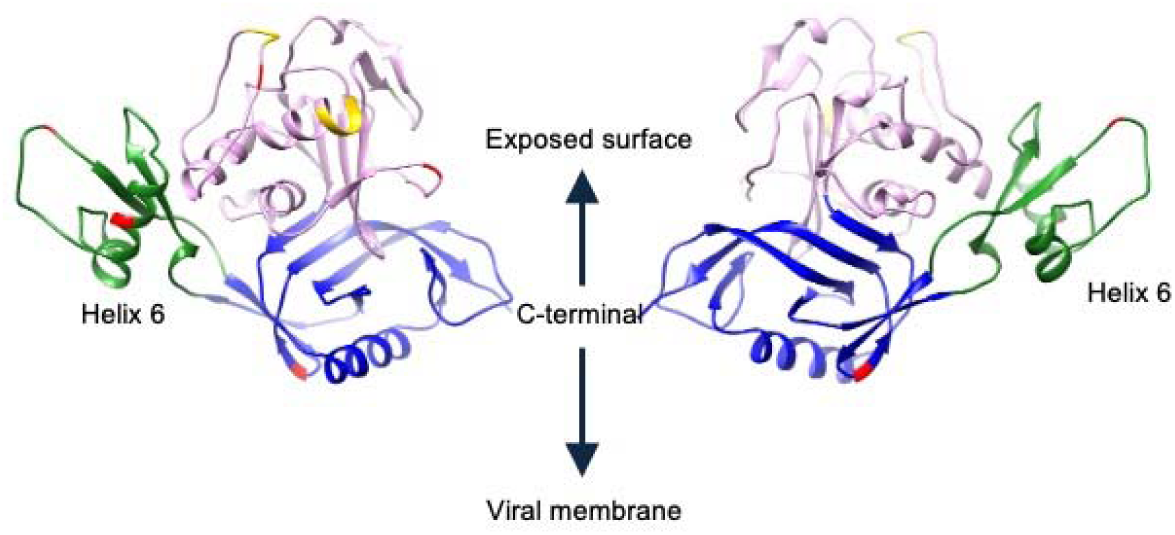
Structural modeling of the Gn glycoprotein head domain. The molecular model highlights the three subdomains of the head region: subdomain I in violet, subdomain II in blue, and subdomain III in green. Amino acid residues that differ between SFTSV strains YL1 and HB29 are indicated in red. Two predicted N-linked glycosylation motifs, NSK33 and NHS63, are shown in yellow and are surface-exposed. Helix 6, a determinant element for neutralizing and viral entry, is indicated. The C-terminal region, from which the stalk and transmembrane domains extend, is also labeled. The model was generated using UCSF Chimera v1.19 based on the crystal structure reported by Wu, Yan, et al. (PDB: 5Y10).(Pettersen et al., 2004; Y. Wu et al., 2017)

**Supplementary Table 1.**
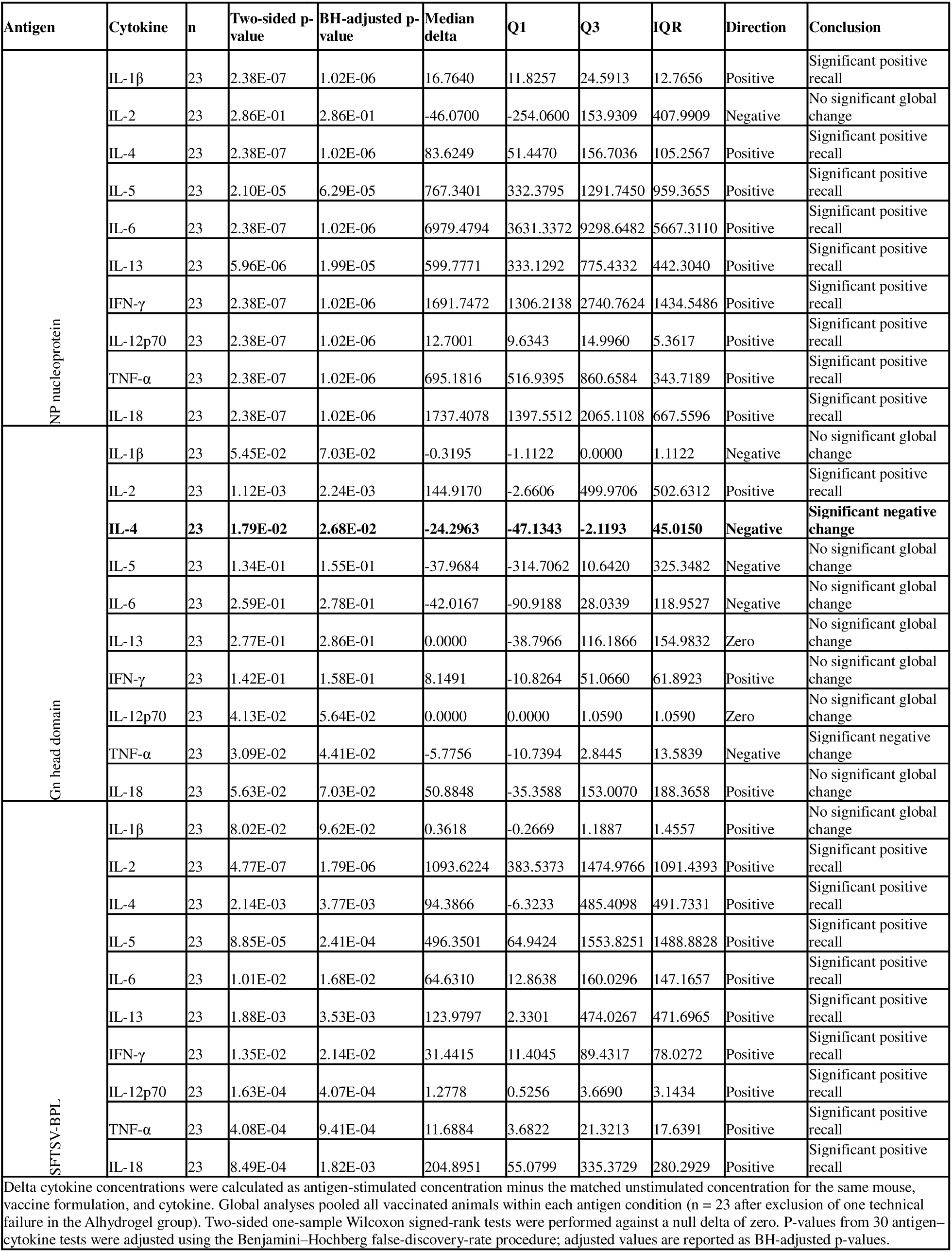
Antigen-Specific Cytokine Recall: Global Analysis.

